# Lactylated histones mark virulence gene families in the malaria parasite *Plasmodium falciparum*

**DOI:** 10.64898/2026.08.12.744368

**Authors:** Ibtissam Jabre, Nana Efua Andoh, Haddijatou Mbye, Alfred Amambua Ngwa, Catherine J. Merrick

**Author notes:** Address correspondence to: Department of Pathology, Cambridge University, Tennis Court Road, Cambridge, CB2 1QP, UK.

## Abstract

Epigenetic pathways have many important roles controlling virulence in human malaria parasites. Histone acetylation and methylation have been closely studied in this context but the novel epigenetic mark of lactylation has not yet been examined. Here, for the first time, we profiled lactyl-histone marks across the *P. falciparum* genome and found them strongly enriched at virulence genes, including genes involved in cytoadhesion and other host-cell remodelling functions. Many genes were dynamically and inducibly lactylated across the cell cycle. Thus, *P. falciparum* could use histone lactylation to control its virulence pathways in response to the prevailing metabolic environment in its host. We extended our chromatin profiling to parasites isolated directly from human patients, showing that here too, virulence gene families were strongly lactylated. This represents the first comprehensive profiling of *P. falciparum* chromatin from parasites in sub-millilitre blood samples, opening up exciting new avenues to study parasite chromatin across human disease states.

## Introduction

*Plasmodium falciparum* is a deeply-diverging protozoan parasite and the most lethal agent of human malaria. Gene expression patterns in this parasite are tightly controlled and much of that control is epigenetic. Epigenetics in *P. falciparum* are particularly influential among families of virulence genes, which can be expressed flexibly with rapid, host-responsive variation [1, 2].

*Plasmodium* parasites have a well-characterised histone code, including modifications like lysine acetylation and methylation that are classically gene-activating or gene-silencing [3]. We recently reported that *Plasmodium* also displays lysine lactylation [4, 5]. This epigenetic mark was discovered in 2019 in human cells [6] and it has since been found in other organisms including several protozoan parasites [7, 8]. The histone code pertaining to this mark is not yet fully characterised.

We hypothesised that histone lactylation could be particularly important in *Plasmodium* biology because malaria characteristically causes hyperlactataemia in human hosts: a clinical predictor of severe disease and fatality [5, 9]. The malaria parasite might therefore have evolved to sense and respond to hyperlactataemia by epigenetically altering its gene expression patterns. Virulence pathways known to be epigenetically controlled, and hence potentially modulated by histone lactylation, include gametocytogenesis (the developmental switch to produce sexual cells adapted for mosquito transmission) [10–12] and the expression of surface adhesins that sequester parasitised cells in the host’s microvasculature [13, 14].

We previously showed that many histone lysine residues in *Plasmodium* are lactylated, inducibly and dynamically, *in vitro* and *in vivo*. Histone lactylation was detected in a range of *Plasmodium* species, including *P. falciparum* (the most important human malaria parasite), *P. knowlesi* (a macaque parasite, zoonotic in humans), and two rodent malaria parasites commonly used as models, *P. yoelii* and *P. berghei*. The species that most characteristically cause hyperlactataemia – *P. falciparum* and *P. yoelii* – displayed the most prominent and dynamic histone lactylation [4]. We also reported that *P. yoelii* grown in hyperlactataemic mice showed major changes in gene expression profiles, and that a selection of *P. yoelii* genes with particularly large expression changes were highly marked with histone lactylation [4]. This implied that the *Plasmodium* transcriptome could be modulated by exposure to high lactate. Nevertheless, the complete landscape of histone lactylation in *Plasmodium*, and its relationship with the transcriptome, was as-yet unexplored.

Here, we focussed on this landscape in *P. falciparum.* We used CUT&Tag, recently adapted for *P. falciparum* [15], to assess the chromatin landscape of *in-vitro*-cultured parasites in parallel with their transcriptome. Then, we harvested parasites directly from human patients in The Gambia (since there is no non-human *in vivo* model for *P. falciparum*) and showed that the signatures detected *in vitro* shared key features with those seen *in vivo*. In particular, the *var* virulence genes that encode cytoadhesion ligands were strongly lactylated both in *vitro* and *in vivo*.

Thus, we present the first landscape of histone lactylation and gene expression for *P. falciparum,* as well as the first proof-of-concept that CUT&Tag chromatin profiling is feasible with *P. falciparum* sampled directly from human blood. Such samples usually have low parasitaemia and high levels of human contamination, making them unsuitable for older techniques like ChIP-seq. This demonstration that they are amenable to CUT&Tag opens up new avenues of epigenetic research in human malaria parasites.

## Results

### CUT&Tag reveals widespread histone lactylation throughout *P. falciparum* chromatin

We initially profiled histone lactylation in the chromatin of trophozoite-stage parasites, because we had previously showed that the highly anabolic trophozoite stage had the highest levels of histone lactylation (KLa), and that some KLa sites were swiftly inducible when trophozoites were exposed to high levels of added lactate [4]. We therefore prepared chromatin from trophozoites under the same conditions as our previous study: exposed or not exposed to 25mM L-lactate over a 12h period.

We selected three KLa antibodies for use in CUT&Tag: a general antibody to lactyl-lysine (‘Pan-KLa’) and two antibodies to specific histone residues, H3K18La and H4K12La. The sequence contexts of both these histone lysines are fully conserved between *P. falciparum* and the human histones whose sequences were used to raise the antibodies [5]. In *P. falciparum,* exposure to exogenous lactate can induce overall lactylation of histones, as detected by the Pan-KLa antibody [4], and H3K18La is a modestly inducible residue (**Fig S1A**), whereas H4K12La is not [4]. We could not survey histones H2A/B, or *Plasmodium*-specific variants like H2A.Z, because these differ from their homologues in humans and suitable antibodies were not available. Nevertheless, the Pan-KLa antibody should detect lactyl-lysine across all histones, as well as other chromatin proteins (many of which are also lactylated [4]). In parallel, we conducted CUT&Tag for acetylated H4: this served to show how acetylation versus lactylation patterns differed.

The CUT&Tag experiments revealed unique distributions of peaks in *P. falciparum* chromatin from all four antibodies (**Fig 1A, S1, Spreadsheet S1**). Lactylation was widely distributed, clearly distinct from H4 acetylation, and not identical between H3K18La, H4K12La and Pan-KLa (**Fig 1A, B**). There were, however, commonalities: telomeric chromatin was highly enriched in all three lactyl marks and this was apparently dependent on the presence of *var* virulence genes. *Var* genes occur in ∼60 hyper-variable copies in *P. falciparum* genomes. They encode an adhesin called *P. falciparum* Erythrocyte Membrane Protein 1 (*Pf*EMP1), which is variantly-expressed on infected erythrocytes. Almost every chromosome has one or more *var* genes in its subtelomeres but chromosome 14 does not, and this chromosome conspicuously lacked subtelomeric KLa peaks (**Fig 1A**). Furthermore, the chromosome-internal tandem arrays of *var* genes were likewise marked with KLa (see chromosome 7, **Fig 1A**).

**Figure 1.**
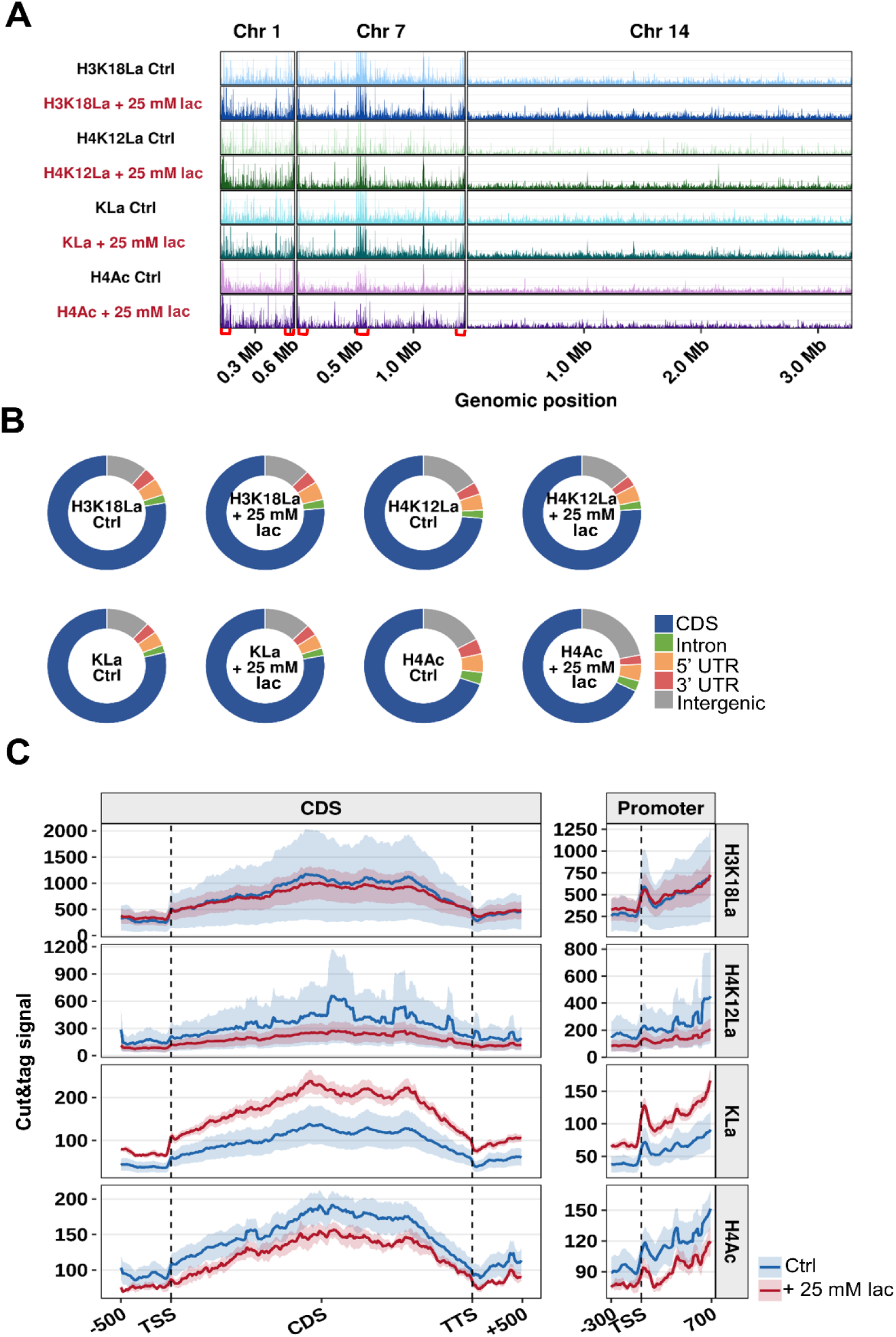
CUT&Tag reveals widespread histone lactylation throughout *P. falciparum* chromatin. **(A)** CUT&Tag profiles of H3K18La, H4K12La, pan-lysine lactylation (KLa) and H4Ac in control parasites (-) or parasites after a 12h exposure to 25mM lactate (+), across 3 representative chromosomes (BigWig signal tracks averaged from 3 biological replicates, track heights scaled to reflect the relative distribution of each modification across the chromosome, rather than absolute signal intensity between histone marks.) *Var* gene arrays on Chr 1 and 7 are indicated by red brackets on the x-axis. **(B)** Genomic annotation of CUT&Tag peaks. Peaks from 3 biological replicates for each histone mark and condition were merged, then assigned according to their greatest overlap with coding sequence (CDS), intron, 5′ untranslated region (5′UTR), 3′ untranslated region (3′UTR) or intergenic sequence. The proportion of peaks assigned to each genomic feature for each histone mark and condition is shown. **(C)** Mean CUT&Tag signals across scaled coding regions and surrounding transcription start sites (TSS). CDS profiles were calculated from 500 bp upstream, across the coding region (scaled to 3kb), plus 500 bp downstream of the transcription termination site (TTS). TSS profiles were calculated from −300 bp to +700 bp using 10-bp bins. Signals were first averaged across genes within each biological replicate and subsequently averaged across replicates. Lines represent mean signal; shading indicates SEM.

Exposure to 25mM lactate, at least over a 12h period, did not greatly change the distribution or number of lactylated peaks (**Fig 1A, Spreadsheet S1**), but many peaks showed changes in intensity. Levels of KLa increased overall (**Fig 1C**), while H4Ac levels, as well as the number of genes detected with H4Ac peaks, reciprocally decreased (**Fig 1C, S1D, E**). This suggested that most of the KLa in *P. falciparum* chromatin exists as a ‘standing’ but tunable mark in the chromatin landscape.

### Histone lactylation is distributed in chromatin differently from acetylation or methylation

Histone lactylation lay primarily within gene bodies (**Fig 1B, C**), distinguishing it from certain important histone acetyl and methyl marks that lie upstream of the genes that they control. The H3K18La signal and the Pan-KLa signal also showed clear peaks at the transcription start site (TSS), whereas these were seen only weakly in H4K12La and H4 acetylation (**Fig 1C**). Histone lactylation evidently appeared as a distinct player in the *P. falciparum* chromatin landscape. To confirm this, we examined its overlap with the known landscapes of euchromatin, heterochromatin, and associated histone marks. Lactylation overlapped strongly with heterochromatin protein 1 (HP1), as expected given its bias towards telomeres and *var* genes (**Fig 2A**), and likewise with a general gene-silencing mark, trimethylated H3K9 (**Fig 2B**). It did not show enriched overlap with trimethylated H3K4, which is generally gene-activating [16] (**Fig 2C**) and it was also anti-correlated with the most accessible chromatin, as detected by ATAC-seq [17] (**Fig 2D**). It showed no correlation with nucleosome density, as defined by micrococcal nuclease sequencing (**Fig 2E**).

**Figure 2.**
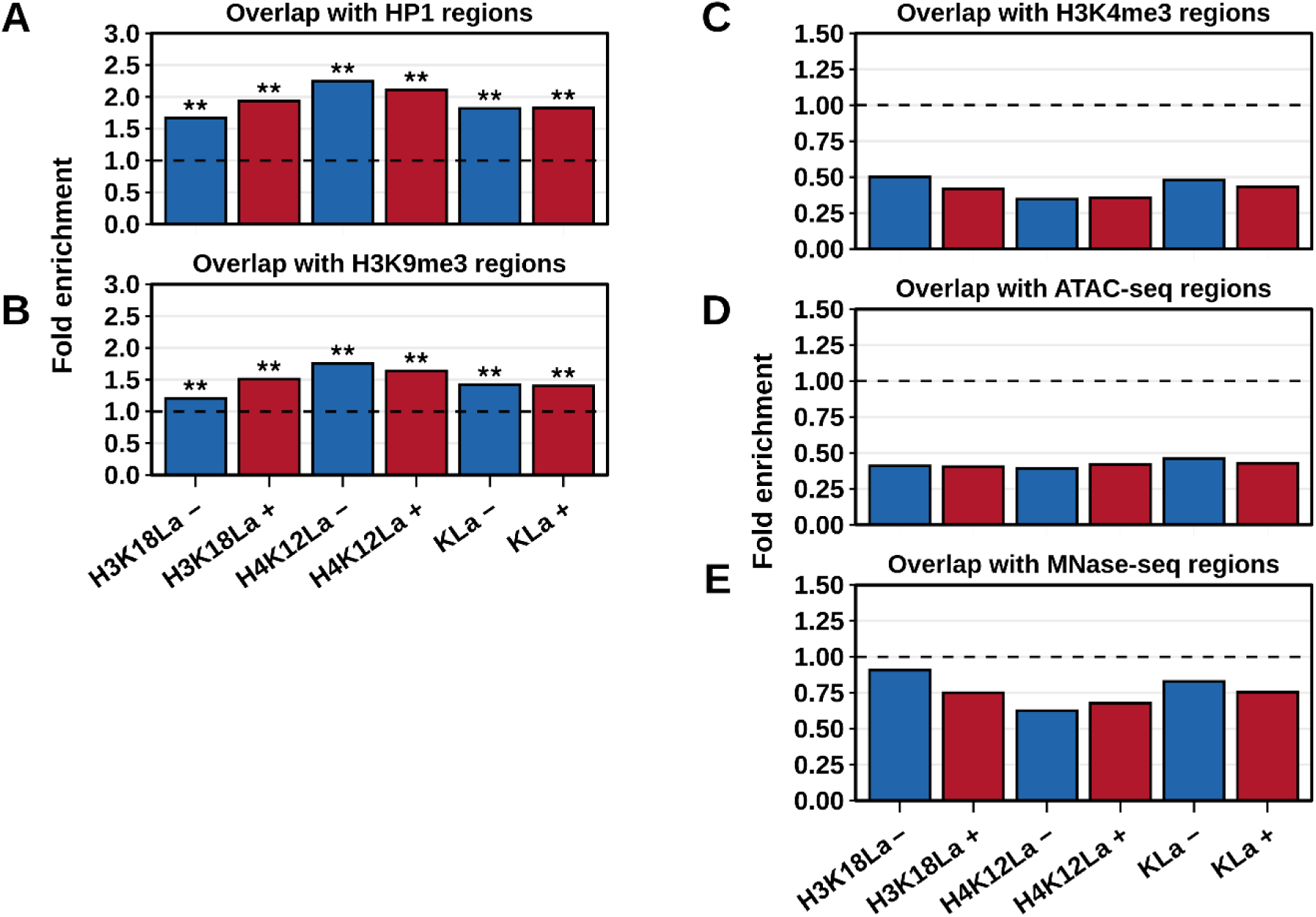
Histone lactylation colocalizes with markers of heterochromatin. Fold enrichment of CUT&Tag peaks overlapping with published chromatin-landscape datasets. Enrichment was calculated as the observed overlap relative to the expected overlap following genomic randomization using the same peak sets. **(A-E)** Fold enrichment of H3K18La, H4K12La and Pan-KLa peaks overlapping HP1-enriched chromatin **(A)**, H3K9me3-enriched regions **(B)**, H3K4me3-enriched regions **(C)**, ATAC-seq-defined accessible chromatin **(D)**, MNase-seq-defined nucleosome-associated regions **(E)**. For all panels, horizontal dashed line (fold enrichment = 1) indicates the overlap expected by random genomic placement. Statistically significant differences from this were determined using permutation-based overlap analysis performed during the original genomic randomization procedure (**P < 0.001, all other comparisons not significant).

### Cytoadhesion genes are amongst the most strongly marked with lactyl histones

Within the landscape of chromatin lactylation, we investigated which gene groups were the most enriched in lactyl-lysine marks. Gene ontology analysis confirmed the picture from Figure 1 that each KLa mark defined a non-identical set of genes, although common GO terms also emerged (**Spreadsheet S2**).

Genes involved in cytoadhesion and host-cell interaction were strongly associated with all KLa marks (**Fig 3A-C**), thus confirming histone lactylation around *var* genes – already apparent from Figure 1. Within these GO terms, only *var*s and the gene for knob-associated histidine-rich protein (KAHRP) were significantly enriched with KLa; other variantly-expressed gene families that cluster together with *var* genes, such as *rifins* and *stevors*, did not reach significant enrichment (**Spreadsheet S2**) Of note, however, KLa and H3K18La were also enriched at *clag* and *Rh* genes, which are not involved in cytoadhesion but are, like *var*s, variantly expressed under epigenetic control [18–20] (**Spreadsheet S2**). H4 acetylation was likewise enriched at *var* genes (**Fig 3D**), since these genes are known to be controlled epigenetically by acetylation versus methylation, but H4KAc was otherwise enriched in very different gene groups from those enriched in KLa.

**Figure 3.**
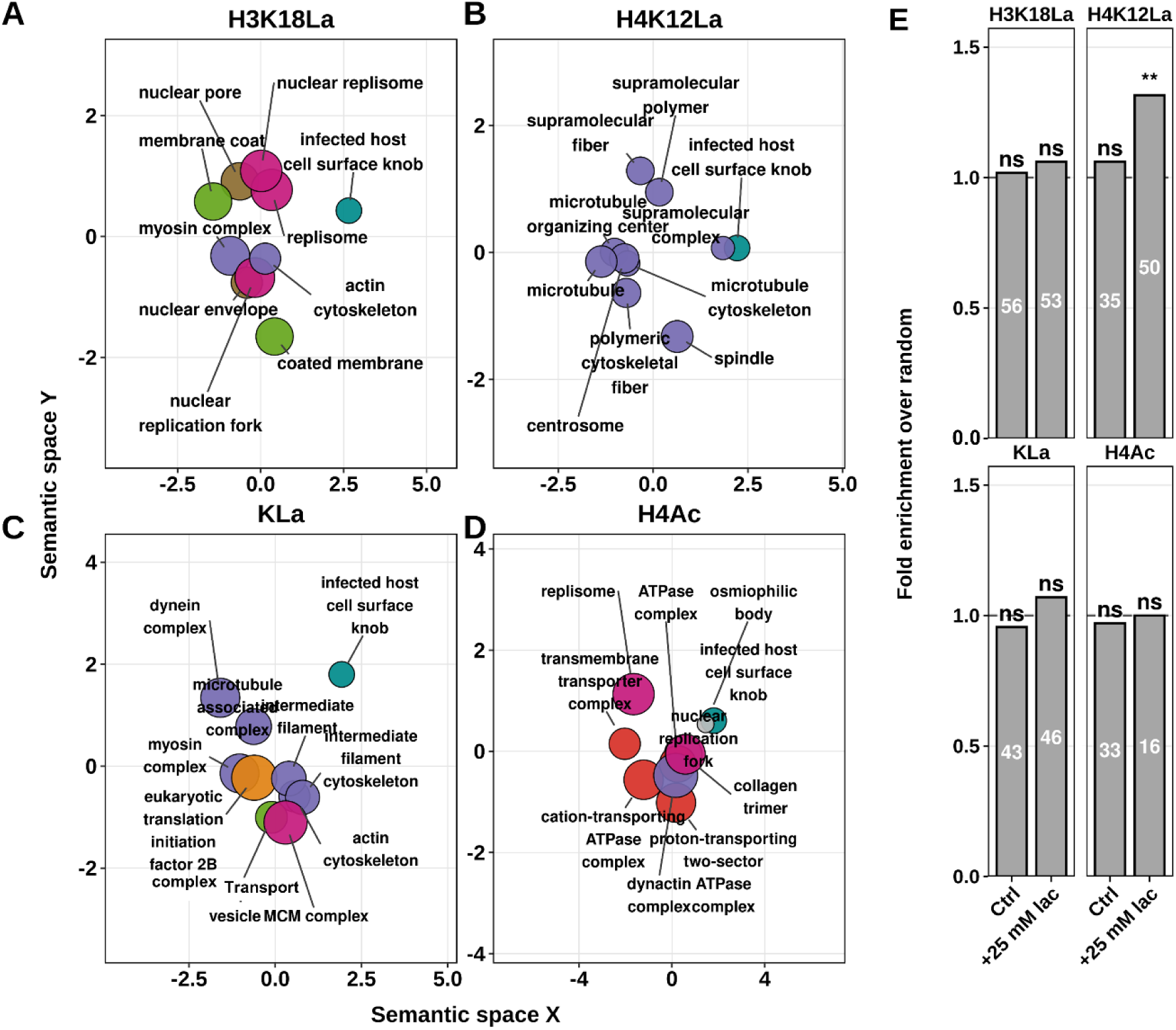
Gene ontology enrichment reveals the gene classes enriched with histone lactylation. **(A–D)** Gene functions associated with CUT&Tag peaks for H3K18La **(A)**, H4K12La **(B)**, pan-lysine lactylation (KLa) **(C)** and H4Ac **(D)**. Gene Ontology (GO) Cellular Component enrichment was performed using genes associated with CUT&Tag peaks for each histone mark. Redundant GO terms were summarized using REVIGO and semantic relationships between GO terms were visualized by classical multidimensional scaling. Bubble size is proportional to GO-term size (LogSize), i.e. the number of genes annotated to each GO term. Bubble outline thickness represents the relative enrichment significance, while bubble colours indicate functionally related GO terms. Labels identify the highest-ranking representative GO terms following semantic redundancy reduction. **(E)** Enrichment of CUT&Tag peaks within gametocytogenesis-associated genes (as annotated on PlasmoDB.org). Fold enrichment was calculated as the observed overlap relative to the overlap expected following genomic randomization. Numbers within the bars indicate the observed numbers of overlapping genes. The horizontal dashed line (fold enrichment = 1) represents the overlap expected by random genomic distribution. Statistical significance was determined by permutation-based genomic randomization analysis (**, adjusted *P* value <0.001; all other comparisons not significant).

Besides the cytoadhesive knobs that contain *Pf*EMP1 and KAHRP, cellular components whose genes were KLa-enriched included the cytoskeleton (enriched for all KLa marks), the phagolysosome, and the replisome and nuclear pore (uniquely for H3K18La) (**Fig 3A-C, Spreadsheet S2**). Among GO terms for biological processes, broadly the same picture was supported: H3K18La marked genes for DNA repair as well as DNA replication; whereas H4K12La marked genes for cytoskeletal components, for phospholipid transport (possibly related to the role of the phagolysosome) and also for RNA capping/processing (**Fig S2**). Finally, we examined genes involved in gametocytogenesis because this is another epigenetically-controlled process influenced by lactate [21] (**Fig 3E**). These genes were specifically enriched with H4K12La in parasites exposed to 25mM lactate; they were not significantly enriched with H4Ac, nor with other KLa marks.

Overall, lactyl-histone marks clearly defined variantly-expressed, epigenetically-controlled genes like *var*s and *clag*s, gametocytogenesis genes, and a wide range of other genes as well. We next set out to establish whether these marks had functional roles in controlling gene expression.

### Exposure of trophozoites to high lactate modulates expression of genes in the exportome and gametocytogenesis pathways

Having established which gene groups showed highly lactylated chromatin, we then proceeded to measure the transcriptomes of the same parasites (i.e. trophozoites exposed or not to high-lactate conditions) via RNA-seq (**Fig S3**), to establish which genes showed acute changes in expression. Hence we could establish whether expression changes correlated with changes in chromatin lactylation.

More than 2000 genes were differentially expressed after 12h exposure to 25mM lactate. Most of the changes were small in magnitude, albeit highly reproducible across replicates (**Fig 4A**): only 49 genes changed expression >2-fold, and 324 genes >1.5-fold (**Fig 4B, Spreadsheet S3**). Most of these genes were up-regulated (42 and 223 respectively) rather than down-regulated (7 and 101 respectively).

**Figure 4.**
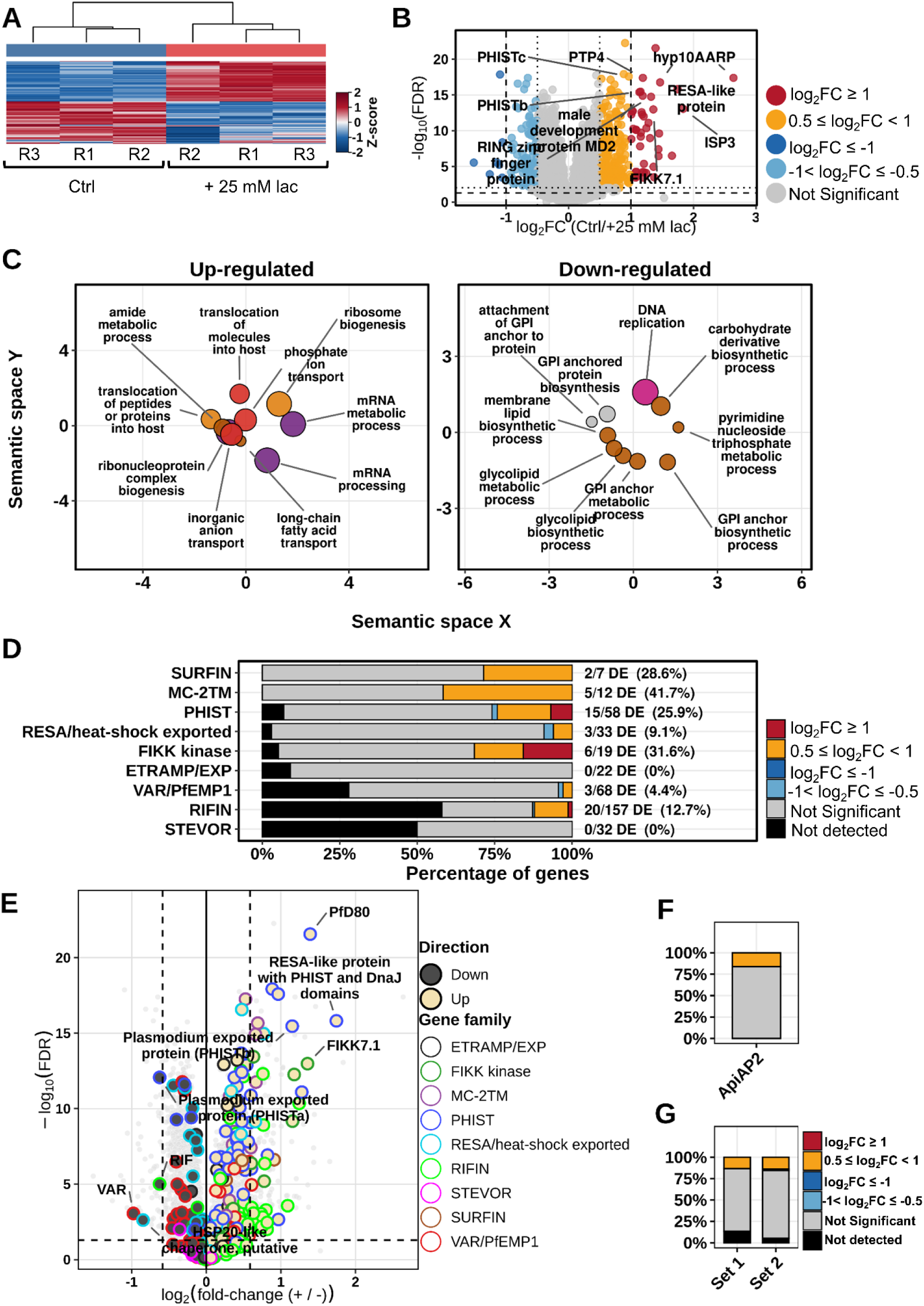
RNA-seq reveals that exposure to high lactate modulates the transcriptome. **(A)** Heatmap showing genes differentially expressed following treatment with 25 mM lactate. Gene expression values were transformed as log₂(expression + 1), standardized by row-wise z-score transformation and hierarchically clustered across both genes and biological replicates. **(B)** Volcano plot summarizing differential gene expression following lactate treatment: Log₂ fold change (25 mM lactate relative to control) plotted versus −log₁₀(FDR). Vertical dashed lines indicate fold-change thresholds of 1.5 and 2-fold. Horizontal dashed and dotted lines indicate the corresponding FDR significance thresholds. Representative significantly regulated genes are labelled. **(C)** REVIGO semantic-space representation of Gene Ontology Biological Process (BP) terms enriched among upregulated and downregulated genes. GO enrichment was performed independently for each gene set, after which redundant GO terms were summarized using REVIGO and plotted as in Figure 3A-D. **(D)** Distribution of differential gene expression across exported and variantly-expressed gene families. Bars show the percentages of genes within each family classified as strongly up/downregulated (≥ 2-fold), moderately regulated or not significantly differentially expressed. The number of differentially expressed genes relative to the total number of annotated genes within each family is shown. **(E)** Volcano plot as in **(B),** highlighting exported and variantly-expressed gene families. Grey points represent all genes analysed by RNA-seq, coloured points identify members of exportome gene families. Selected significantly regulated genes are annotated. **(F)** Differential expression of ApiAP2 transcription factors following 12 h exposure to 25 mM lactate. The bar shows the proportion of ApiAP2 genes classified according to differential-expression category, as in **(D)**. **(G)** Differential expression of gametocytogenesis-associated genes, plotted as in **(F)**, shown separately for a small subset of regulatory genes (set 1) and for the complete gametocytogenesis-associated gene set annotated on PlasmoDB.org (set 2).

Gene ontology analysis showed that downregulated genes had GO terms for growth, such as DNA replication and biosynthetic processes, whereas many upregulated genes were involved in RNA processing, or were components of the exportome (**Fig 4C**). Large numbers of *SURFIN, PHIST, RESA, PfMC-2TM* and *FIKK* genes were detected (**Fig 4D, Spreadsheet S3**), many with >1.5-fold changes (**Fig 4D, E**). A smaller proportion of *var* genes were differentially expressed, probably because their expression occurs in the ring stage of cellular development, whereas these transcriptomes were from more mature trophozoite stages, in which *var* genes are largely silenced whereas most of the other exportome families are expressed.

The main family of *Plasmodium-*specific transcription factors is termed Api-AP2 [22], and these are master regulators for developmental programmes, including production of invasive merozoites [23] and production of sexual transmission stages [10–12]. Genes in the Api-AP2 family were almost universally upregulated after lactate exposure, albeit modestly, with most showing fold-changes under the 1.5 threshold (**Fig 4F**). Gametocytogenesis genes were also upregulated, including *AP2-G, AP2-G2*, *AP2-G5,* the male development regulator *Md2* and the upstream regulator of *AP2-G*, *GDV-1* (**Fig 4G, Spreadsheet S3**). Changes were again modest (∼1.25 to 1.5-fold), but importantly only a minority of parasites in any population usually convert to sexual stages within a single growth cycle, thus diluting their expression signature. We hypothesised that a subset of early regulatory genes like *GDV-1* and *AP2-G* might show greater changes than the whole set of >60 genes associated with gametocytogenesis, but this was not the case (**Fig 4G**, set 1 vz set 2).

### Histone lactylation can be either activating or silencing in different gene groups

We sought concordance between differentially-expressed genes and differential lactylation in the chromatin of those genes. The majority of genes were not directly concordant (**Fig 5A**), probably because acute changes in gene expression are primarily controlled as whole pathways via lactate-induced transcription factors, rather than via direct changes to histone lactylation at every gene. Nevertheless, several hundred genes did show concordance between expression and chromatin levels of lactylated histones, whereas very few genes showed such concordance with H4Ac (**Fig 5A**). (Changes at H2A/B, which we did not measure, could also be involved, but these should be encompassed in the Pan-KLa CUT&Tag dataset). There was no evidence that specific UTR-based marks particularly influenced gene expression: changes in lactylation were mostly restricted to the CDS, wherein most of the KLa marks reside (**Fig 5B**).

**Figure 5.**
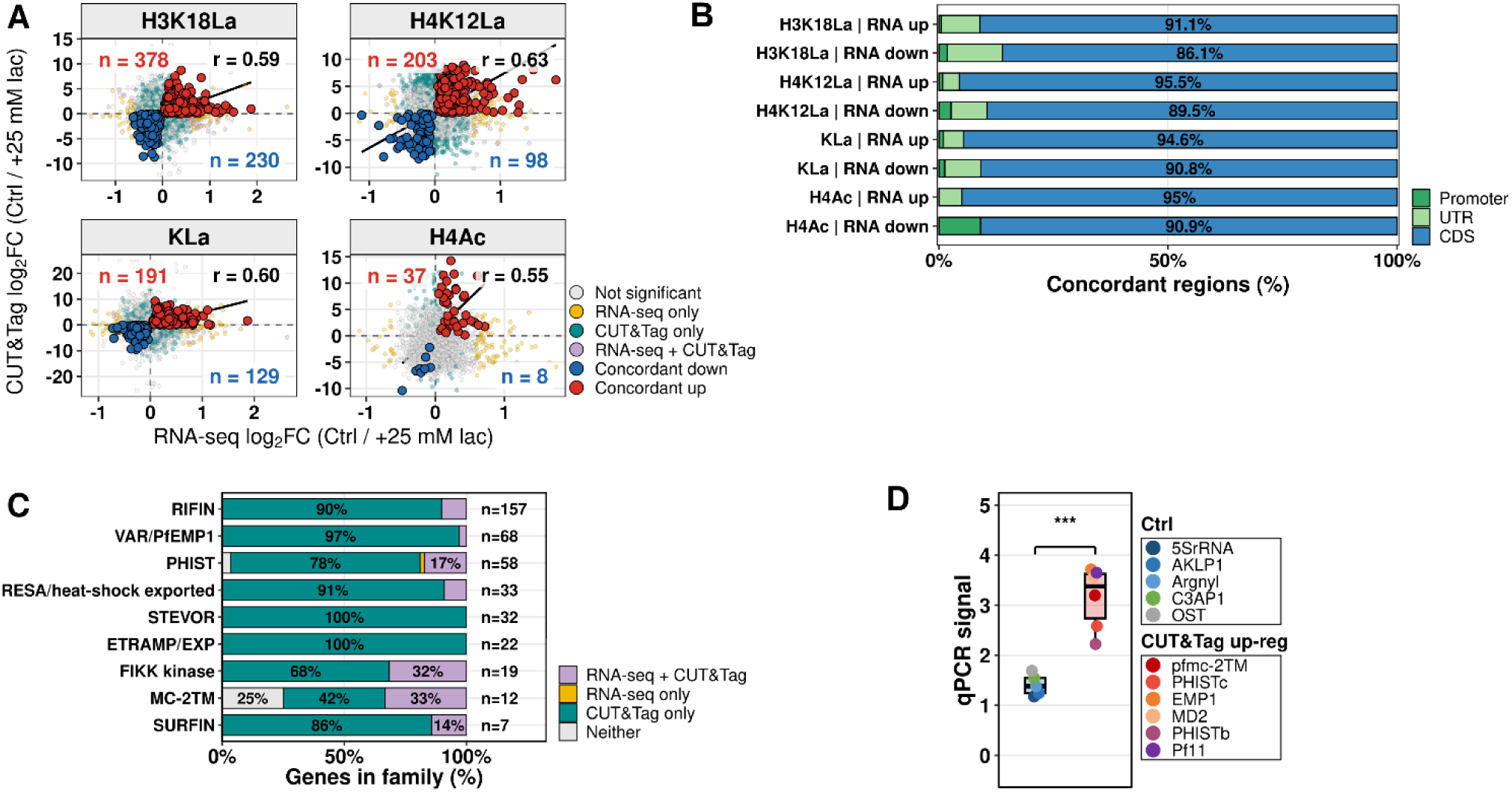
Integration of CUT&Tag and RNA-seq identifies coordinated epigenetic and transcriptional responses to lactate. **(A)** Relationship between transcriptional and chromatin changes following lactate treatment. Scatter plots show differential expression in RNA-seq (x-axis) and CUT&Tag log₂ fold changes (y-axes) for H3K18La, H4K12La, pan-lysine lactylation (KLa) and H4Ac. Genes were classified according to the significance thresholds applied independently to the RNA-seq and CUT&Tag analyses (RNA-seq only, CUT&Tag only, significant in both datasets, or non-significant). Concordantly upregulated and concordantly downregulated genes are highlighted and their numbers are shown. The black line represents a linear regression fitted for visualization, and the reported correlation coefficient corresponds to Spearman’s rank correlation. **(B)** Genic distributions of CUT&Tag regions concordantly associated with transcriptionally regulated genes. Concordant regions were assigned to the promoter, untranslated region (UTR) or coding sequence (CDS) according to the genomic feature with which each region showed the greatest overlap. Bars show the percentage of concordant regions assigned to each genomic feature and percentages were calculated relative to the total number of concordant regions within each comparison. **(C)** Integration of RNA-seq and CUT&Tag data across exported and variantly-expressed gene families. Genes were classified according to whether significant changes were detected by RNA-seq, CUT&Tag, both datasets or neither. Bars show the proportion of genes belonging to each category within individual families and numbers indicate the total number of genes per family. **(D)** Experimental validation of selected genes by ChIP-qPCR using the H3K18La antibody. Six genes were selected that showed concordant changes in RNAseq and Cut&Tag, versus a control gene group that did not. Box plots represent the distribution of ChIP-qPCR ratios obtained for each gene in parasites treated with 25mM lactate, versus untreated. The mean ratio within each group is also shown. Statistical significance was assessed using a two-sided Wilcoxon rank-sum test comparing gene-level mean expression between the two groups.

Changes in the chromatin landscape of differentially-expressed genes occurred in both directions – up- and down-regulation – and in both directions the magnitudes of chromatin changes and expression changes tended to correlate (**Fig 5A**). This implies a complex histone code in which lactyl marks are associated with activation in some genes but repression in others. Gene ontology analysis supported this (**Fig S4A, Spreadsheet S4**): for example, genes involved in DNA replication and repair were concordantly downregulated in expression and in H3K18La, whereas genes involved in the cytoskeleton were upregulated in expression and lactylation. Focussing on the exportome genes that were dominantly upregulated in expression (Figure 4), a proportion of those genes also showed significant changes in their CUT&Tag signals, reaching up to ∼30% of the *FIKK* and *MC-2TM* families (**Fig 5C**). Most such changes were positively concordant, i.e. expression and chromatin lactylation were both induced, but a smaller number of genes involved in knobs showed concordance in the opposite direction (**Fig S4A**) – a pattern that might indicate lactate-associated ‘switching’.

Finally, we validated our data on a small selection of genes by conducting new, independent lactate-exposure experiments, followed by gene-by-gene chromatin immunoprecipitation and qPCR. This confirmed that the CUT&Tag data were robust and reproducible: genes upregulated in their CUT&Tag signal were also upregulated in their ChIP signal after lactate exposure, whereas a group of control genes were not (**Fig 5D, S4B**).

### *Var* genes have distinctive, promoter-specific patterns of histone lactylation

The chromatin of *var* genes was strongly lactylated (**Fig 1**), yet their expression changed relatively little in the RNA-seq dataset (**Fig 4D**), probably because *var* genes are epigenetically silenced in the trophozoite stage. They are, however, epigenetically ‘poised’, with a histone-code memory that marks the previously-expressed gene for re-expression in the next cell cycle [24].

A detailed examination of the chromatin lactylation signature of the *var* gene family revealed a unique enrichment of H3K18La, and to a lesser extent H4K12La, ∼100bp upstream of the CDS (**Fig 6A**). Such marking was not present upstream of non-*var* genes. It appeared very similar to the known enrichment of H4KAc in *var* gene promoters (**Fig 6A**). It was not strongly affected by 12h of high-lactate exposure in trophozoites, suggesting a ‘standing’ chromatin signature of H3K18La and H4K12La, but the Pan-KLa antibody did detect inducibility, presumably on other histone residues or other chromatin proteins (**Fig 6A**).

**Figure 6.**
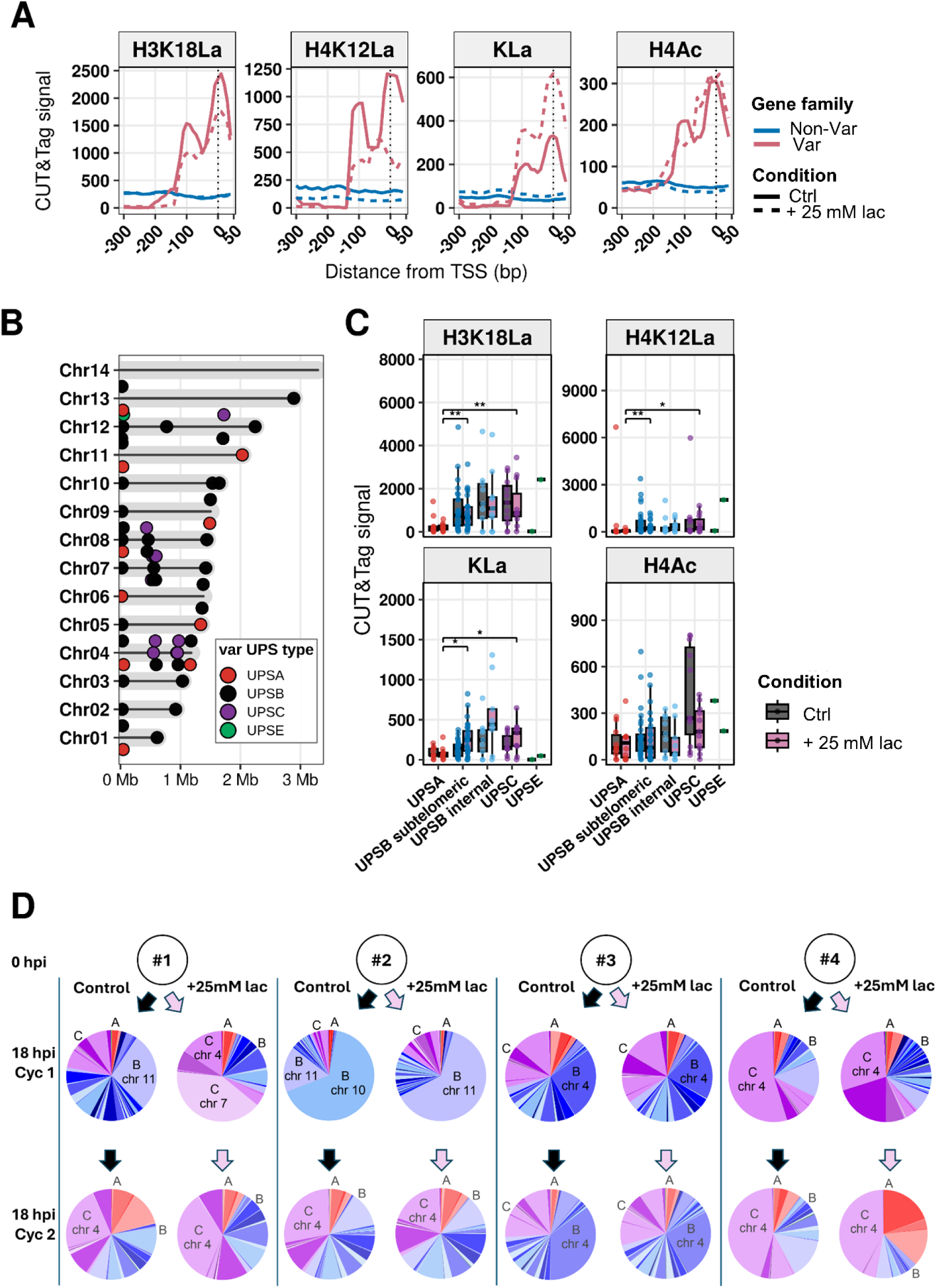
Histone lactylation patterns in *var* virulence genes are distinctive. **(A)** Metagene profiles for the promoters of *var* and non-*var* genes. CUT&Tag signal was calculated from 300 bp upstream to 50 bp downstream of the TSS using 10-bp bins, then averaged across all genes within each group and subsequently across biological replicates. Lines represent the mean CUT&Tag signal for H3K18La, H4K12La, KLa and H4Ac under control and 25 mM lactate conditions. The vertical dashed line denotes the TSS. **(B)** Chromosomal distribution of *var* genes colour-coded according to upstream sequence (UPS) class. Each point represents the midpoint of an individual *var* gene; points are vertically offset where necessary to improve visualization. **(C)** CUT&Tag signal upstream of upsA, upsC, upsE, subtelomeric upsB and chromosome-internal upsB *var* genes. The mean CUT&Tag signal from 300 bp upstream to 50 bp downstream of the TSS was calculated for each gene, shown as boxplots under control and 25 mM lactate conditions. Statistical comparisons between control and lactate-treated parasites were performed independently for each histone modification and promoter class using two-sided Wilcoxon rank-sum tests with Benjamini–Hochberg correction for multiple testing. **(D)** Relative proportions of *var* genes expressed across the gene family in 4 independent clones of NF54. Ups groups are colour-coded (red, upsA; blue, upsB; purple, upsC; green, upsE) and the ups-type and chromosomal location of the dominant *var* transcript at each timepoint is labelled. *Var* expression was assessed in synchronised populations of ring stages at∼18hpi after 18h of culture with or without 25mM lactate, and again after a further full cycle of growth and reinvasion in the same conditions.

*Var* genes can be subdivided into upsA, B, C and E types according to their promoter type and chromosomal position (**Fig 6B**) [25]. Lactylation of the 5’ UTR was strongest on upsC and weakest on upsA genes (**Fig 6C**). Where a gene type could be either subtelomeric or chromosome-internal (upsB or ‘B/C’ genes), lactylation was strongest in the chromosome-internal group (**Fig 6C**). The upsE gene, which is subtelomeric but has a unique structure and involvement in placental sequestration during pregnancy malaria [26], showed a notable absence of lactylation in its 5’ UTR, but this was highly inducible in high-lactate conditions, which reciprocally reduced the upsE gene’s acetylation (**Fig 6C**).

We investigated whether these ups-specific patterns of lactylation might influence which type of *var* gene is actually expressed in ring-stage parasites, using an established panel of q-RTPCR primers to assess *var* expression patterns across the entire *var* gene family in several clones of the NF54 strain [26, 27]. Each clone was exposed or not exposed to 25mM lactate from 0 to 18hpi (the point in the ring stage when *var* expression peaks), and RNA was harvested for *var* expression profiling. Then both cultures were continued in the same conditions through reinvasion to the next ring stage for a second RNA harvest.

This revealed that the established tendency of cultured parasites to switch towards, and then stably express, upsC genes [28] may actually be promoted by lactate (of which significant levels are produced by high-density cultures [4]) (**Fig 6D**). Three out of four clones began by dominantly expressing three different subtelomeric upsB genes, and two of these readily switched to one or more upsC genes. In clone #1, the switch was precipitated within 18h of exposure to added lactate, whereas switching to upsC occurred in the control culture only after a full cycle of growth and reinvasion, with associated endogenous production of lactate. In clone #2, lactate rapidly induced an initial switch to another upsB gene, then after 1 cycle the culture switched to a dominant upsC gene, along with a diversity of minor upsB genes (again, this same switch towards upsC occurred after 1 growth cycle without 25mM added lactate). Clone #3, by contrast, expressed a different upsB gene and appeared ‘fixed’: it did not switch regardless of lactate exposure. Finally, clone #4 already expressed a upsC gene and within 18h it showed no ‘’switch’, only some diversification among other chromosome-4 upsC genes. Interestingly, after another cycle, the same upsC gene remained dominant but the lactate-treated population also expressed several upsA genes: the only case among these four clones where large amounts of upsA expression occurred.

### Lactylation of *var* genes occurs *in vivo* as well as *in vitro*

We next investigated whether our *in vitro* data accurately represented *P. falciparum* biology *in vivo.* There is no accessible non-human model for *P. falciparum* malaria, so parasites must be sampled directly from human patients. Only ring-stages can be sampled from peripheral blood because trophozoites and schizonts are largely sequestered in the microvasculature via their cytoadhesive knobs.

Parasites in the low-volume blood samples routinely taken in malaria clinics tend to be sparse and highly contaminated with human leukocytes. We therefore developed a protocol for removing the leukocytes and preparing the parasites for CUT&Tag, exploiting the low-input, low-noise, high-sensitivity nature of this chromatin profiling technique (**Fig S5A**). We demonstrated for the first time that CUT&Tag could be successfully conducted on less than 1ml of blood at sub-1% parasitaemia, and thus obtained data on the genome-wide distribution of H3K18La from 6 samples (**Fig S5B-G**). We focussed here on H3K18La as the mark with the strongest signature in *var* gene chromatin. In parallel, we conducted CUT&Tag on ring-stage NF54 parasites cultured *in vitro*, since these should match the staging of the field-isolated parasites better than the existing trophozoite-stage datasets (**Fig S5, Spreadsheet S5)**.

The chromatin of field isolates mirrored cultured parasites remarkably closely. *Var* genes stood out as being strongly lactylated in all samples, so this signature evidently exists in ring stages as well as trophozoites (**Fig 7A**). GO terms associated with cytoadhesion and antigenic variation were accordingly the most enriched among lactylated genes in all 6 field isolates (**Fig 7B, S6A**). Other GO terms matching the *in vitro* data also appeared: for example, genes associated with the phagolysosome were enriched in H3K18La in all of the field isolates. Other gene groups, however, such as those for DNA replication and the cytoskeleton, were not strongly enriched (**Fig 7B, S6A, Spreadsheet S6**).

**Figure 7.**
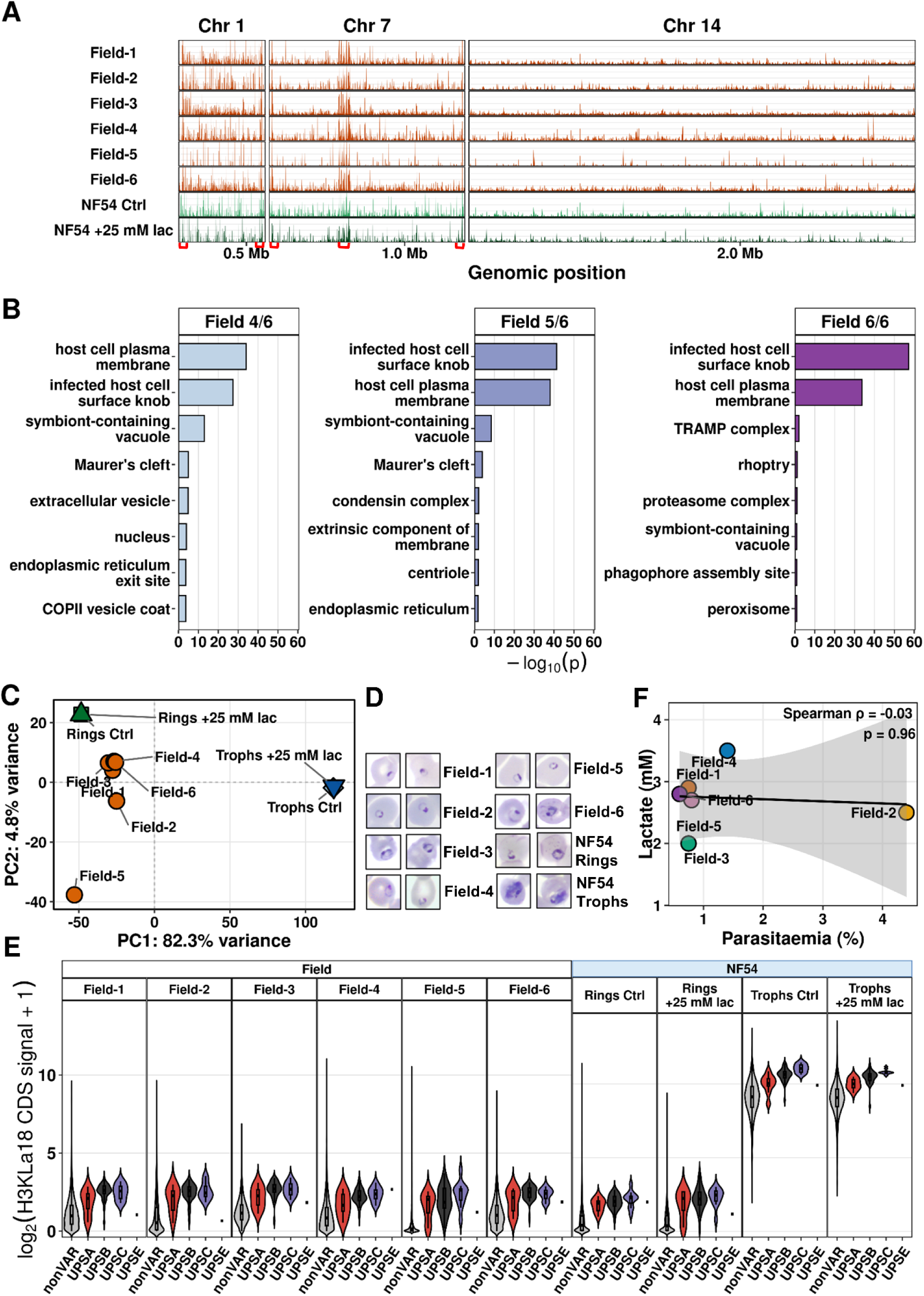
Lactylation of *var* virulence genes is strong and distinctive *in vivo* as well as *in vitro*. **(A)** Genome-wide distribution of H3K18La across representative chromosomes (1,7 and 14) in field isolates and laboratory-cultured NF54 ring-stage parasites, plotted as in Figure 1A. Signals from 3 biological replicates were averaged for NF54 ring-stage parasites, whereas each field isolate was analysed by Cut&Tag once and is shown individually. Var gene arrays on Chr 1 and 7 are indicated by red brackets on the x-axis. **(B)** Gene Ontology Cellular Component (CC) enrichment of genes associated with H3K18La peaks shared between *P. falciparum* field isolates. Genes were grouped according to the reproducibility of peak association across the six field isolates (genes in at least four, five or six out of six). Cellular Component GO enrichment was performed separately for each group. **(C)** Principal component analysis (PCA) of H3K18La signal across the genomes of field isolates, NF54 rings and NF54 trophozoites. NF54 biological replicates were averaged within each condition before PCA. Signals were transformed as log₂(signal + 1), centred and scaled prior to analysis. Axes indicate the percentage of variance explained by the first two principal components. **(D)** Representative Giemsa-stained blood smears showing morphology of parasites used in all CUT&Tag experiments. **(E)** Distribution of H3K18La signal in CDS across non-*var* genes and *var* genes with different promoter subclasses in field isolates, NF54 rings and NF54 trophozoites. The y-axis shows gene-level mean log₂(CUT&Tag signal + 1). Violin plots represent the distribution of genes within each category, while embedded boxplots indicate the median and interquartile range. **(F)** Relationship between peripheral parasitaemia and circulating lactate concentration in the clinical blood samples. Linear regression was fitted for visualization, correlation coefficient corresponds to Spearman’s rank correlation.

In contrast to the chromatin from NF54 trophozoites, chromatin from field isolates additionally showed H3K18La enrichment in non-*var* exportome families like *rifin, stevor* and *PHIST.* GO terms such as ‘Maurer’s cleft’ were enriched here, but not in the NF54 trophozoite dataset. Thus, the H3K18La mark on these gene families appears to be stage-specific in ring stages (**Fig 7B, S6A, Spreadsheet S6**), whereas marking of replication and cytoskeleton genes appears to be stage-specific in trophozoites. (Stage-specificity was further investigated in the following section.)

Figure 7a revealed that despite their overall similarities, H3K18La chromatin profiles did vary between field isolates. Principal component analysis (PCA) (**Fig 7C**) showed that all isolates clustered in the dominant component, together with ring-stage NF54 and away from the trophozoites, so this component clearly separated the samples according to cell-cycle stage. The minor component, however, showed more variation – not apparently stage-related because parasites that appeared as more mature rings (e.g. isolates #3 and #6, **Fig 7D**) did not cluster away from the others. The minor component may be related to density of lactylation, since some isolates showed much more than others (compare, for example, all non-*var* genes in isolates #5 versus #6, **Fig 7E**).

When *var* genes were specifically examined, their patterns of lactylation mirrored the patterns in cultured NF54 rings (**Fig 7E**): *var* genes were always strongly lactylated (both within (**Fig 7E**) and upstream of the CDS (**Fig S6B**), although their marking was still less abundant in rings than in trophozoites. Furthermore, NF54 rings showed variation in *var* lactylation depending on whether or not they had been exposed to 25mM lactate (**Fig 7E**), and in the field isolates, *var* gene lactylation generally resembled that seen in NF54 rings and most closely resembled NF54 rings exposed to high lactate (**Fig 7E**).

These patterns could, speculatively, indicate that the field isolates were exposed to high lactate, or possessed an epigenetic memory of earlier lactate exposure *in vivo*. This pilot study was not, however, designed to address the question of epigenetic memory because samples were drawn at only one timepoint. Blood lactate and parasitaemia were measured at this timepoint, but the cohort was very small and quite homogenous. At the time of sampling, all six blood samples fell below the clinical threshold for hyperlactataemia (2-3.5mM), and parasitaemias were, with one marked exception, around 1%: there was no correlation between host lactataemia and parasitaemia (**Fig 7F**).

### Chromatin lactylation occurs stage-specifically across the erythrocytic cycle

We observed in Figure 7E that the epigenetic landscape of histone lactylation was much stronger in trophozoites than ring stages. Since trophozoites are the most metabolically active stage, producing the most lactate and having the most strongly lactylated histones overall [4], this was perhaps unsurprising. Nevertheless, there could be stage-specificity not only in the overall abundance of the modification but also in its detailed distribution across the genome. Therefore, we compared the landscapes of KLa and H3K18La in NF54 rings (∼18hpi) versus trophozoites (∼36hpi).

The two stages correlated, but only moderately (R ∼0.4 in gene CDSs and ∼0.25 in upstream regions (**Fig 8A, S7A**)), so the lactylation of many genes did indeed change with stage. The GO terms most enriched in ring stages still included those associated with knobs on the host cell membrane (hence *Pf*EMP1), as well as terms associated with the phagolysosome. However, terms for DNA replication and the cytoskeleton were missing from the top-10 (as they were in field-isolated ring stages), whereas terms for protein production and cell growth, such as ‘ribosome’ and ‘endoplasmic reticulum’, appeared in the top-10 (**Fig 8B, S7B**).

**Fig 8:**
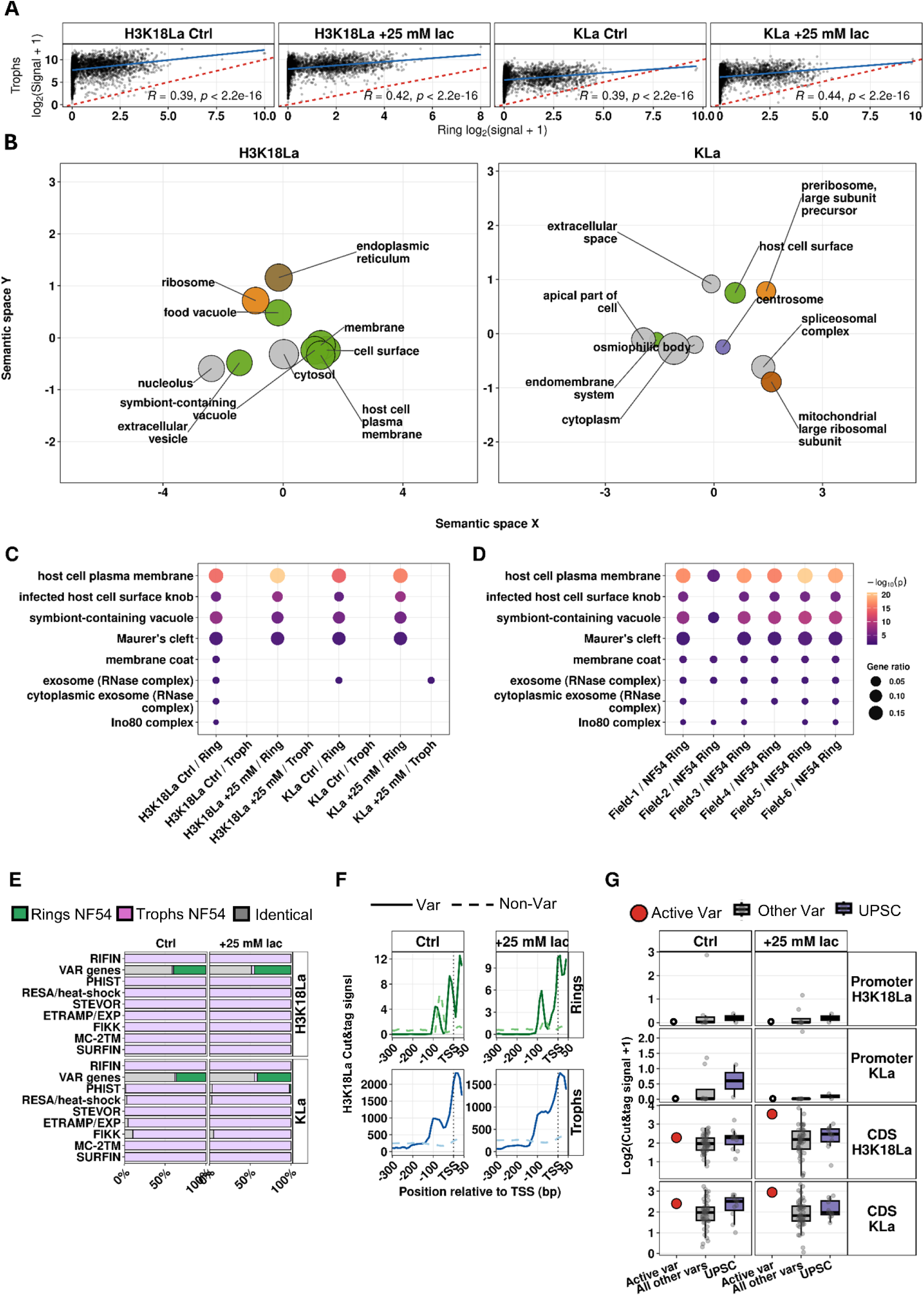
Stage-specificity of chromatin lactylation patterns. **(A)** Gene-level comparison of H3K18La and pan-lactylation (KLa) CUT&Tag signal over coding sequences in NF54 ring- and trophozoite-stage parasites under control and 25 mM lactate conditions. Each point represents one gene and signals are shown as log₂(signal + 1). The red dashed line indicates equal signal between stages and the blue line shows the fitted linear regression. Spearman correlation coefficients and associated *P* values are shown within each panel. **(B)** REVIGO semantic-space representation of Cellular Component Gene Ontology terms associated with the union of genes linked to H3K18La or Pan-KLa peaks detected in NF54 ring-stage parasites across control and 25 mM lactate conditions. The ten most significant terms are shown for each mark. Spatial proximity reflects semantic similarity between GO terms. Bubble size represents GO-term size, bubble-outline thickness indicates enrichment significance, and bubble colour denotes related functional term families. **(C)** Cellular Component GO enrichment of genes showing relative H3K18La or KLa enrichment between NF54 ring- and trophozoite-stage parasites. Gene-level CUT&Tag signal was quantified over common strand-aware coding-region intervals (+50 to +1,050 bp relative to the transcription start site), transformed as log₂(signal + 1), and independently robust z-score normalised for each stage, mark and treatment condition. Relative stage scores were calculated as the ring-stage robust z-score minus the trophozoite robust z-score. Genes in the upper and lower deciles of each distribution were classified as relatively ring-enriched and trophozoite-enriched, respectively, and Cellular Component GO enrichment was performed separately for each gene set. Dot size represents gene ratio and dot colour represents enrichment significance (−log₁₀ *P*). **(D)** Cellular Component GO enrichment of genes showing relatively stronger H3K18La enrichment in NF54 ring-stage parasites than in individual field isolates. Each of the six field isolates was compared independently with the averaged NF54 ring-control reference. Gene-level signal was quantified over the same coding-region intervals used in (C), log₂-transformed and independently robust z-score normalised. Relative scores were calculated as the field-isolate robust z-score minus the NF54 ring-control robust z-score, and genes in the lower decile, corresponding to relative enrichment in NF54 rings, were used for GO enrichment. For (C) and (D), Cellular Component over-representation analysis was performed using the same *P. falciparum* GO background; the strongest eight terms from each comparison were considered and, for visualisation, only terms represented in at least three comparisons were retained. This final filter affected display only and did not alter the underlying enrichment analyses. Dot size represents gene ratio and dot colour represents −log₁₀ *P*. **(E)** Relative stage enrichment across individual exported-protein gene families and *var* genes. Bars show the percentage of genes classified as ring-enriched, trophozoite-enriched or showing no strong stage difference for each family, separately for H3K18La and KLa under control and 25 mM lactate conditions. **(F)** Metagene profiles of H3K18La CUT&Tag signal across *var* and non-*var* genes in rings and trophozoites under control and 25 mM lactate conditions, plotted as in Figure 6A. Ring-stage and trophozoite-stage profiles are shown separately for *var* and non-*var* genes. **(G)** Distribution of H3K18La and Pan-KLa CUT&Tag signal among the active *var* gene, all other *var* genes and the UPSC *var* subgroup. Data are shown separately for promoter and coding-sequence regions under control and 25 mM lactate conditions. Boxes indicate the median and interquartile range, whiskers show the distribution outside the interquartile range, and individual genes are displayed as points. Signals are expressed as log2 (CUT&Tag signal + 1). Signals for upsC versus other *var* genes were compared using two-sided Wilcoxon rank-sum tests with Benjamini–Hochberg correction for multiple comparisons; no comparisons were significant after adjustment.

This suggested that histone lactylation could be dynamic across the cell cycle, with genes becoming marked at the stage of their peak transcriptional activity. Indeed, genes related to knobs and Maurer’s clefts were over-enriched in lactylation in ring stages compared to trophozoites (**Fig 8C**). (Interestingly, they were also relatively *more* enriched in cultured NF54 rings than in most of the field isolates (**Fig 8D**), with isolate #2 – which had the highest parasitaemia (**Fig 7F**) – being the exception.) By contrast, exportome genes as a whole were less lactylated at the ring stage. Figures 4-5 showed that in trophozoites, many exportome gene families were strongly upregulated after lactate exposure, and some of these genes were concordantly lactylated. In ring stages, however, only the *var* genes were more strongly marked whereas all other exportome families were more strongly marked in trophozoites (**Fig 8E**). Again, this may relate to the cell-cycle stage at which each gene is most transcriptionally active. Specifically at *var* genes, we checked the unique upstream peak of lactylation that was seen in trophozoites (**Fig 6A**) and although the peak still existed (**Fig 8F**), lactyl-lysine abundance was many-fold lower at these loci, as it was overall.

Finally, we examined the chromatin of the *var* gene that was actively transcribed. This Cut&Tag experiment used Clone#1 from Figure 6D, which started out expressing a upsB gene and switched under high lactate exposure to upsC. Strikingly, the upstream region of this active gene was completely without detectable KLa, whereas the upstream regions of other genes – notably upsC genes – were lowly but detectably marked. Conversely, within the CDS, all *var* genes were marked at similar levels and high lactate caused a particular elevation at the active gene. Once again, this suggests that active transcription allows chromatin to become highly lactylated – and perhaps (at least among *var* genes) this then contributes to silencing of expression and hence a ‘switch’.

## Discussion

This study provides the first genome-wide view of histone lactylation in *P. falciparum.* It also provides the first evidence that this low-abundance epigenetic mark can be effectively profiled using Cut&Tag, not only in cultured parasites but in field isolates from human patients. This allows us to make unprecedently direct comparisons of epigenetic signatures *in vitro* and *in vivo*, which is particularly illuminating for histone lactylation given its possible role as a host-parasite signal [5]. Importantly, this Cut&Tag protocol opens up the general prospect of studying malaria parasite chromatin in the immediate context of human disease states.

We found that *P. falciparum* chromatin was broadly lactylated across many genes, most notably *var* and cytoadhesion-related genes such as *KAHRP*, but also genes associated with DNA replication and repair, the cytoskeleton, and the food vacuole, amongst others. The functional significance of all these modules being regulated via lactate is yet to be determined, but they may contribute to a metabolic shift designed to downregulate growth in high-lactate conditions, as we saw in *P. yoelii* exposed to hyperlactataemia [4]. Besides cytoadhesion genes, which we had previously linked to host lactate levels in a study of human patient isolates [29], we also examined genes involved in gametocytogenesis because this developmental switch was recently linked to lactate via *in vitro* experiments [21]. Indeed, some gametocytogenesis genes were enriched in lactylation when parasites were exposed to high lactate. Thus, at least three virulence phenotypes – cytoadhesion, gametocytogenesis and growth rate – may be regulated by lactate.

Chromatin lactylation did not, overall, move dramatically after an acute 12h exposure to exogenous lactate, suggesting that it is a ‘standing’ but also ‘tunable’ mark in the epigenetic landscape. A large number of genes did change in expression in response to lactate, probably because whole expression pathways are up- or downregulated by lactate-induced regulators like ApiAP2 transcription factors. Overall, transcripts were primarily upregulated rather than downregulated, many *AP2* genes were modestly upregulated, and gametocytogenesis genes, which are controlled by a module of AP2 factors [10–12], were, as expected, upregulated. In addition, there was a striking upregulation of the exportome: *PHIST*s, *FIKK*s, etc. Hyperlactataemia may therefore be sensed by *P. falciparum* as a signal to upregulate all aspects of host cell remodelling, as well as to upregulate the host-exit strategy of gametocytogenesis. Interestingly, febrile temperatures can also stimulate cytoadhesion via increased trafficking through the exportome [30]. The parasite may have evolved to sense and respond similarly to two different signals of host stress: hyperlactataemia and fever [29].

This picture in *P. falciparum* differs somewhat from what we previously observed in the transcriptome of *P. yoelii* [4]. Those *P. yoelii* parasites had grown for much longer in hosts developing hyperlactataemia, alongside other aspects of pathology, and their transcriptomes changed greatly, with a broad metabolic shift downregulating replication and glycolysis and showing signatures of ‘dormancy’ [4]. Here, we applied the high-lactate stimulus in isolation for just a brief 12h period, and *P. falciparum* did not show a similar metabolic shift within 12h. We hypothesise, however, that longer lactate exposures would downregulate replication in *P. falciparum* as well, because at the molecular level, many replication genes were regulated by lactylation, while at the phenotypic level, we observed that the cell cycle was increasingly delayed at the point of reinvasion if parasites were lactate-exposed for more than 1 cell cycle ([4] and observations during this study).

The lactylation code is evidently complex. Altered gene expression after lactate exposure correlated with either increased or decreased chromatin lactylation, depending on the gene type. This may be determined by the ‘code’ – the exact combination or location of lactyl marks – or it may be determined by the presence of proteins that write and erase histone lactylation. These have yet to be identified in *Plasmodium*, but by analogy with human cells they probably include ‘moonlighting’ histone acetyltransferases and deacetylases (HATs and HDACs) [5], so the lactyl code probably acts in concert with the acetyl code.

The lactyl code is not only complex, it is evidently also dynamic with respect to the cell cycle. When comparing ring and trophozoite stages, we observed that the genes most enriched with lactylation tended to be those actively transcribed at that stage (such as genes for DNA replication in the trophozoite stage). Active transcription may render a gene available for lactylation, and transcription-coupled writer enzymes may then respond to the prevailing levels of lactate in the cell. This would explain exactly how this metabolic signalling axis works, but further experiments are needed to validate such a hypothesis.

We focussed our further analysis upon how the lactyl code imposes upregulation of exportome genes in response to lactate. Most exportome genes were lactylated primarily within their CDS, and when lactylation was induced, expression was usually elevated in the trophozoite stage. Thus, in most of these gene families, lactylation correlates with chromatin opening and active transcription (either by inducing it, or by *resulting from* it). *Var* genes, by contrast, which are silenced in the trophozoite stage, showed a unique lactylation peak upstream of the CDS (similar to the acetylation that occurs in the same region). In ring stages, where a single *var* gene is active, this was maintained but at vastly lower levels, and the active gene showed none at all, whereas that gene was still strongly and inducibly marked in its CDS. Therefore, lactylation upstream of *var* loci may have a unique role in gene silencing, and this may be inducible by high lactate exposure. By contrast, lactylation within the CDS may build up as the active gene is transcribed, and then perhaps promote switching if it eventually spreads upstream. *Var* expression switching is already known to be mediated by stochastic epigenetic changes in acetylation versus methylation. We observed that high lactate exposure tended to promote switching away from upsB towards upsC genes, and that the CDS of an active (upsB) gene was particularly lactylated, commensurate with this model.

How this might affect upsA and upsE genes remain unclear: lactylation was almost absent upstream of upsA genes, but these are rarely strongly activated *in vitro* [28]. The upsE gene has a unique ‘node’ role in *var* gene switching [31] and interestingly its upstream region was unlactylated, but highly inducible by exogenous lactate. This might affect the unique uORF encoded there [31]: a topic for further study.

The data reported herein compare interestingly with our earlier study of virulence gene expression in parasite isolates from humans, which was conducted long before histone lactylation was discovered [29]. This focussed on *var* genes and sirtuin HDACs, and revealed a correlation between sirtuin expression, upsB *var* expression and host hyperlactataemia. Parasites in a ‘high-sirtuin/high-upsB’ state should cytoadhere strongly, promoting hyperlactataemia, whereas upsC expression is associated with milder malaria [32–34], so it could be advantageous for parasites to sense the lactate signal and downregulate a host-endangering state by switching from upsB to upsC expression. To find out how sirtuins might be involved in this, their possible role as histone delactylases must be elucidated in future work.

Meanwhile, to truly establish cause and effect between hyperlactylation and switching, a longitudinal study would be needed with repeated chromatin profiling across many different *var-*expressing clones. This would be feasible, if laborious, *in vitro*, but difficult to conduct in a truly relevant disease setting, i.e. untreated human malaria patients. Furthermore, it is an intriguing possibility that spikes of hyperlactataemia, experienced *in vivo* by parasites during previous growth cycles, could remain as ‘epigenetic memory’ to regulate *var* gene expression in subsequent cycles. Longitudinal studies would again be very difficult to conduct, particularly *in vivo*, but some of these questions may be addressed correlatively, using large cohorts of patient samples taken at varying blood lactate levels. Our demonstration herein that Cut&Tag is feasible in field samples – the first ever such Cut&Tag study in *P. falciparum*, to our knowledge – makes such studies a realistic and exciting prospect for the future.

## Materials and Methods

### Parasite culture, synchronization and lactate treatment

*Plasmodium falciparum* NF54 parasites (BEI Resources, www.beiresources.org) were maintained in human O⁺ erythrocytes at 4% haematocrit in RPMI 1640 (Sigma, R4130) supplemented with 25 mM HEPES, 2.3 g L⁻¹ sodium bicarbonate, 50 mg L⁻¹ hypoxanthine (Sigma, H9377), 25 μg mL⁻¹ gentamicin (Melford Laboratories, G38000-1), 2.5 g L⁻¹ Albumax II (Invitrogen, 11021037) and 5% human serum. Cultures were maintained at 37°C under a gas mixture of 3% O₂, 5% CO₂ and 92% N₂. Parasite growth, developmental stage and parasitaemia were monitored routinely by microscopic examination of thin blood smears.

Synchronization was as in Jabre *et al.* [4] and independent experiments were performed using synchronized trophozoite- and ring-stage parasites. For trophozoite-stage experiments, synchronized cultures were allowed to develop to ∼24–26 h post-invasion (hpi) and adjusted to 5% parasitaemia. Immediately before treatment, existing medium was replaced to minimise variation resulting from lactate accumulation during parasite growth. Cultures were divided into matched control and treatment groups derived from the same synchronized population. Sodium L-lactate (Sigma, L7022) was freshly prepared in incomplete RPMI medium and added at a final concentration of 25 mM, while control cultures received an equivalent volume of incomplete medium without lactate. Parasites were maintained under these conditions for 12 h before collection for western blotting, CUT&Tag, RNA sequencing or ChIP–qPCR.

Ring-stage experiments were performed using tightly synchronized parasites treated as above 2 h after erythrocyte invasion. Parasites were incubated for a further 18 h before harvesting for CUT&Tag analysis.

Unless otherwise stated, all experiments were performed using three independent biological replicates. Control and lactate-treated cultures were established from the same synchronized parasite population and processed simultaneously throughout each experiment.

### Western blotting

Histone H3 lysine 18 lactylation (H3K18La) was quantified by western blotting using synchronized trophozoite-stage parasites cultured in the absence or presence of 25 mM sodium L-lactate for 12 h. Three independent biological replicates were analysed for each condition.

Following treatment, infected erythrocytes were lysed in 0.1% (w/v) saponin prepared in ice-cold phosphate-buffered saline (PBS), and released parasites were recovered by centrifugation. Parasite pellets were washed thoroughly in ice-cold PBS before extraction in RIPA buffer (250 mM Tris-Hcl pH 8.0, 750 mM NaCl, 5% NP-40, 5% sodium deoxycholate, 0.5% SDS, 1mM phenylmethylsulfonyl fluoride) containing 1X cComplete EDTA-free protease inhibitor cocktail (Roche, 0469313200). Protein extracts were generated by repeated freeze–thaw disruption, mixed with Laemmli loading buffer and denatured at 95°C. Protein concentrations were determined before electrophoresis, for loading of 30 μg total protein per lane, and western blotting was conducted as in [4]. Band intensities were quantified using Fiji-win64 (Image J). H3K18La signal intensity was normalized to the corresponding H4 signal for each biological replicate, and normalized values were expressed relative to the untreated control.

### CUT&Tag

Genome-wide profiling of histone lactylation and histone acetylation was performed using Cleavage Under Targets and Tagmentation (CUT&Tag), adapted from the protocol described by Gockel et al. [15].

Libraries were generated from tightly synchronized NF54 trophozoite- and ring-stage parasites cultured under control conditions or following treatment with 25 mM sodium L-lactate. Independent libraries were prepared for H3K18La, H4K12La, pan-lysine lactylation (KLa) and H4 acetylation (H4Ac), with three biological replicates generated for each histone modification and treatment condition. Parallel IgG controls were processed throughout the workflow and subsequently used as the matched background during peak identification.

Infected erythrocytes were lysed in ice-cold 0.1% (w/v) saponin prepared in PBS. Released parasites were washed extensively in PBS supplemented with freshly prepared EDTA-free protease inhibitor cocktail (Roche) before fixation in 1% methanol-free formaldehyde (Thermo Scientific, 28906) for 8 min at 37°C with gentle agitation (250 rpm). Crosslinking was terminated by addition of glycine to a final concentration of 125 mM, after which parasites were washed in ice-cold PBS and immediately processed for nuclei isolation.

Parasite nuclei were isolated using sequential hypotonic lysis followed by permeabilization in cell lysis buffer (.10 mM Tris-HCl (pH 8.0), 3 mM MgCl₂, 0.2% v/v Nonidet P-40, 0.25 M sucrose, and a cocktail of EDTA-free protease inhibitors (Roche, 0469313200)) Approximately 3.5 × 10⁷ trophozoite nuclei or 5–10 × 10⁶ ring-stage nuclei were resuspended in Cut&Tag wash buffer (20 mM HEPES pH 7.5, 150 mM NaCl, 0.5 mM Spermidine, 0.1% Triton X-100, 1× Protease Inhibitor Cocktail) per reaction. Purified nuclei were immobilized on activated Concanavalin A magnetic beads (Bangs Laboratories, BP531).

Bead-bound nuclei were incubated overnight at 4°C with each antibody (concentrations optimized during method development and maintained throughout the study) resuspended in antibody buffer (0.5 M EDTA, 10%BSA in Cut&Tag wash buffer). Parallel IgG reactions were processed under identical conditions. Following primary antibody incubation, nuclei were washed once with CUT&Tag wash buffer (20 mM HEPES pH 7.5, 150 mM NaCl, 0.5 mM Spermidine, 0.1% Triton X-100, 1× Protease Inhibitor Cocktail) and incubated for 1 h at room temperature with the corresponding secondary antibody, then washed three times in CUT&Tag wash buffer and once with CUT&Tag-300 wash buffer (20 mM HEPES pH 7.5, 300 mM NaCl, 0.5 mM Spermidine, 0.01% Digitonin, 1× Protease Inhibitor Cocktail), before binding of CUTANA™ protein A/G-Tn5 transposase (EpiCypher, 15-1017) for 2 h.

Following removal of excess transposase by washing the nuclei one time with CUT&Tag-300 wash buffer, tagmentation was initiated by addition of CUT&Tag 300-Wash Buffer supplemented with MgCl_2_, and carried out for 1 h at 37°C. Tagmentation was stopped by addition of EDTA (43.9 mM), SDS (0.26%, w/v), and Proteinase K (0.44 mg mL⁻¹) at 55°C for 1 h to digest proteins and release tagmented DNA. DNA was purified using the Genomic DNA Clean & Concentrator-10 kit (Zymo Research; D4011). Overnight incubation at −20°C following addition of DNA-binding buffer consistently improved DNA recovery, particularly from ring-stage samples, and this step was therefore incorporated into all experiments. DNA was recovered by silica-column purification and eluted twice using pre-warmed elution buffer to maximise yield. DNA concentration was determined using a Qubit High Sensitivity DNA assay before library preparation (Invitrogen, Q32851).

Sequencing libraries were prepared as described previously [15]. Final libraries were quantified using KAPA Library quantification Kit (Roche, 7960417001) and their fragment-size distributions were assessed using TapeStation prior to pooling.Libraries were sequenced on an Illumina platform using paired-end 100-cycle (P2) chemistry. Approximately 400 million paired-end reads were generated across the complete CUT&Tag dataset.

### Preparation of patient isolates for CUT&Tag

Patients with uncomplicated *Plasmodium falciparum* malaria were recruited at Brikama Health Centre, The Gambia. For each patient reporting with fever and suspicion of malaria, *Plasmodium spp* infection was diagnosed by RDT, followed by microscopy. All those characterised with uncomplicated malaria were invited to participate.

Following written informed consent, ∼2 mL of peripheral venous blood was collected into EDTA tubes, which were immediately stored and transported in cooler boxes (∼4°C) to the laboratories at the Medical Research Council Unit The Gambia at the London School of Hygiene & Tropical Medicine (MRCG at LSHTM) for processing on the day of collection. For minors (>18 years), signed consent was obtained from a parent or legal guardian, with assent obtained where applicable. Ethical approval for the parent study was obtained from The Gambia Government/MRC Joint Ethics Committee (Ref: SCC 1626v1.2).

Blood lactate was measured using a Lactate Pro 2 meter (Arkray), then ∼1mL of whole blood was washed in ice-cold RPMI-HEPES medium, diluted 10x in ice-cold RPMI-HEPES, and depleted of host leukocytes using Plasmodipur filtration (EuroProxima, 8011). Filtered erythrocytes were collected by centrifugation, parasites were released by saponin lysis and washed extensively in PBS containing protease inhibitors before formaldehyde fixation as above. Prior to nuclei isolation, fixed parasites were frozen at −80°C and stored for international shipping.

Subsequent processing of patient isolates followed the same workflow established for cultured parasites, including nuclei isolation, Concanavalin A bead immobilisation, antibody incubation, pA/G-Tn5-mediated tagmentation, DNA purification and library preparation. H3K18La was profiled in individual field isolates, with each isolate processed and sequenced as an independent biological sample.

### CUT&Tag data analysis

Raw CUT&Tag sequencing reads were subjected to quality assessment before downstream analysis. Adapter sequences and low-quality bases were removed where necessary prior to alignment against the *Plasmodium falciparum* 3D7 reference genome using STAR (v2.7.11b). Alignments were performed independently for each biological replicate, and coordinate-sorted BAM files were generated for all libraries. Reads mapping to the apicoplast or mitochondrial genomes, together with low-quality alignments where present, were excluded before downstream analyses. Genome-wide coverage tracks were generated from the filtered and normalized alignments and exported in BigWig format for quantitative analysis and visualization. To reduce the impact of differences in sequencing depth between field isolates, libraries with substantially higher read depth were randomly downsampled before generation of normalized BigWig files. Specifically, isolates #3, 5 and 6 were downsampled to approximate the sequencing depth of the lower-coverage isolates #2 and 4, which were analysed without modification. The resulting depth-matched BigWig files were used for genome-wide comparative analyses of the field isolates, including principal component analysis and gene-level signal quantification. This normalization was performed solely to minimise sequencing-depth bias and did not alter peak calling or other analyses performed using the original alignments. Biological replicates were retained independently throughout quality control and statistical analyses and were combined only where indicated for graphical presentation.

Regions of significant enrichment were identified independently for each biological replicate using **SEACR** (version 1.3) with the corresponding IgG library used as the matched background control. Consensus peak sets were generated separately for each histone modification and treatment condition by merging overlapping peaks identified across biological replicates, thereby generating a single non-redundant peak set for downstream analyses.

For CUT&Tag quality-control analyses, fraction of reads in peaks (FRiP) was calculated directly from sample-matched BAM files and SEACR peak sets. Properly paired primary fragments were extracted using SAMtools, excluding unmapped, mate-unmapped, secondary, supplementary, duplicate and quality-control-failing alignments. Fragment coordinates were reconstructed from paired-end alignments and intersected with the corresponding sample-specific SEACR peaks using bedtools intersect -u. FRiP was calculated as the number of usable mapped fragments overlapping at least one peak divided by the total number of usable mapped fragments. GC-content bias was assessed across the 14 nuclear chromosomes of the PlasmoDB annotation release 82, *P. falciparum* 3D7 reference genome. The genome was divided into complete non-overlapping 300-bp windows and the GC fraction of each window was calculated from the reference sequence. For each CUT&Tag BAM, properly paired primary alignments were counted as fragments and the number overlapping each genomic window was determined using GenomicRanges::countOverlaps(), excluding unmapped, secondary, supplementary, duplicate and quality-control-failing alignments. Windows were grouped into GC-fraction bins of width 0.01 and local fragment depth was plotted against GC content to assess GC-dependent sequencing bias across trophozoite, NF54 ring-stage and field-isolate CUT&Tag libraries.

Genomic annotation was performed relative to the *P. falciparum* 3D7 reference annotation (PlasmoDB/VEuPathDB release 82) by reconstructing strand-specific transcript models from annotated transcript, exon and coding-sequence coordinates. Coding sequences (CDS), 5′ untranslated regions (5′UTRs), 3′ untranslated regions (3′UTRs) and introns were reconstructed directly from the genome annotation, while genomic intervals outside annotated transcripts were classified as intergenic. Where a consensus peak overlapped multiple genomic features, the peak was assigned to the feature sharing the largest base-pair overlap, ensuring that each peak contributed to a single genomic category during downstream analyses.

Genome-wide CUT&Tag signal was quantified directly from normalized BigWig files. For chromosome-scale visualization, each chromosome was divided into approximately 1,200 equally spaced genomic bins, and the mean signal intensity was calculated for each bin. To facilitate comparison of signal distribution along individual chromosomes, each track was independently truncated at its 99.5th percentile and scaled between zero and one before visualization. This normalization was applied exclusively for graphical representation and was not used for statistical comparisons.

Gene-level CUT&Tag signal was quantified independently across the genomic interval relevant to each analysis. TSS-proximal signal was calculated across the fixed promoter interval extending from 300 bp upstream to 50 bp downstream of the annotated TSS. Where this was compared directly with untranslated regions, reconstructed annotated 5′UTRs were analysed independently.

Chromatin signals were subsequently integrated with RNA-seq datasets using common gene identifiers derived from the *P. falciparum* reference annotation. RNA-seq and CUT&Tag results were analysed independently before integration. Genes were classified according to whether statistically significant changes were detected by RNA-seq only, CUT&Tag only, both datasets or neither dataset. Concordant gene sets comprised genes exhibiting significant chromatin and transcriptional changes in the same direction and formed the basis of the integrated analyses presented throughout the manuscript.

Chromatin enrichment with each of the 4 epigenetic marks analysed was further quantified across curated biological gene sets obtained from their annotation on PlasmoDB.org (ref), including exported protein families, ApiAP2 transcription factors, gametocytogenesis-associated genes and variant (*var)* genes. *Var* genes were classified according to their annotated upstream sequence (UPS) class. Chromosomal distributions of *var* genes were generated using the annotated chromosome lengths. Promoter and coding-sequence enrichment were analysed independently for each UPS class. UPSB genes were further subdivided into subtelomeric and internal subclasses according to their chromosomal location.

To compare developmental stages and field isolates while accounting for global differences in CUT&Tag signal intensity, Cellular Component enrichment across cultured and field-isolate CUT&Tag datasets was assessed using gene-level signal quantified over common strand-aware coding-region intervals extending from +50 to +1,050 bp relative to the transcription start site. For NF54 stage comparisons, log₂(signal + 1) values were independently robust z-score normalised within ring- and trophozoite-stage datasets for each antibody and treatment condition, and a relative stage score was calculated as the ring-stage robust z-score minus the trophozoite robust z-score. Genes in the upper and lower deciles of this distribution were classified as ring-enriched and trophozoite-enriched, respectively. For field comparisons, mean raw H3K18La signal across the three NF54 ring-control replicates was calculated for each gene and subsequently log₂-transformed, whereas each field isolate was analysed independently using its log₂-transformed gene-level signal. Field and NF54 ring-control distributions were independently robust z-score normalised, and the relative score was calculated as the field-isolate robust z-score minus the NF54 ring-control robust z-score. Genes in the lower decile, representing relative enrichment in NF54 rings, were retained for the final field-isolate comparisons. Cellular Component over-representation analysis was performed separately for every gene set using clusterProfiler and the PlasmoDB-68 *P. falciparum* 3D7 GO annotation, with all Cellular Component-annotated genes used as the background, Benjamini–Hochberg correction, a minimum term size of three genes and a maximum term size of 5,000 genes. The strongest eight terms from each comparison were considered for visualisation, and the final matrix was restricted to terms occurring in at least three comparisons to reduce sparse rows; this final restriction affected display only and did not alter the underlying enrichment analyses. Dot size represents gene ratio and dot colour represents −log₁₀(P).

### RNA sequencing

Total RNA was isolated from synchronized trophozoite-stage *Plasmodium falciparum* NF54 parasites maintained under control conditions or exposed to 25 mM sodium L-lactate for 12 h. Three independent biological replicates were prepared for each condition. Control and lactate-treated samples were generated from the same synchronized parasite population and processed in parallel throughout RNA extraction and library preparation.

Following treatment, infected erythrocytes were immediately resuspended in TRIzol Reagent (Thermo Fisher Scientific, Cat. No. 15596026). Chloroform was added at 0.2 volumes, and samples were centrifuged at 9,000 × g for 30 min at 4°C to separate the aqueous phase. Total RNA was purified from the aqueous phase using the RNA Clean & Concentrator-5 kit (Zymo Research, Cat. No. R1015) according to the manufacturer’s instructions. Residual genomic DNA was removed by on-column DNase I digestion, and purified RNA was eluted in nuclease-free water supplemented with SUPERase•In™ RNase Inhibitor (Invitrogen, Cat. No. AM2694).

RNA concentration was determined using NanoDrop and RNA integrity was assessed using an Agilent Bioanalyzer RNA 6000 Nano assay. Only high-quality RNA preparations were used for downstream analyses. The absence of detectable genomic DNA contamination was confirmed by PCR [28] using the following cycling conditions: 95°C for 2 min, followed by 40 cycles of 95°C for 5 s and 60°C for 30 s.

Polyadenylated mRNA was enriched from total RNA using the NEBNext Poly(A) mRNA Magnetic Isolation Module (New England Biolabs, Cat. No. E7490), employing two rounds of oligo(dT)-based selection. First-strand cDNA synthesis was performed using SuperScript III RNase H-Reverse Transcriptase (Thermo Fisher Scientific, Cat. No. 18080044), followed by second-strand synthesis to generate double-stranded cDNA. Library preparation was based on the principles of the DAFT-seq protocol for AT-rich genomes [35], in which full-length cDNA is generated prior to fragmentation to improve transcript coverage across the highly AT-rich *P. falciparum* genome.

Instead of the original DAFT-seq workflow, double-stranded cDNA was processed using the NEBNext Ultra II DNA PCR-Free Library Preparation Kit (New England Biolabs, Cat. No. E7410), including end repair, A-tailing, adapter ligation and library cleanup according to the manufacturer’s recommendations. Unique dual-index adapters were ligated to each library, which was prepared without PCR amplification to minimise sequence-composition bias. Final libraries were quantified using KAPA Library quantification Kit (Roche, 7960417001) and their fragment-size distributions were assessed using TapeStation prior to pooling.

Equimolar libraries were sequenced on an Illumina platform using paired-end 100-bp reads. Approximately 100 million paired-end reads were generated across the complete RNA-seq dataset.

### RNA-seq data analysis

RNA-seq data were processed using the analytical framework applied in [4], omitting stage deconvolution analysis. In brief, raw sequence quality was assessed using FastQC (v. 0.11.9). Adapter sequences and low-quality bases were removed using Trimmomatic v.0.35; bases with Phred quality scores below 30 were trimmed and reads shorter than 20 nucleotides after trimming were discarded. Filtered reads were quantified against the *P. falciparum* 3D7 reference transcriptome (PlasmoDB/VEuPathDB release 82) using Salmon v.0.82 with sequence-bias correction and bootstrap sampling enabled. Transcript-level abundance estimates were imported into R using tximport and summarized at gene level using length-scaled transcript-per-million estimates.

Genes that reached at least 1 count per million in a minimum of three libraries were retained; those with insufficient expression were removed before differential-expression modelling. Library composition was normalized using the trimmed mean of M-values method. Mean–variance relationships were modelled using voom, and differential expression between control and 25 mM lactate-treated trophozoites was assessed using the limma linear-modelling framework with empirical Bayes moderation. The model contrasted lactate-treated parasites directly with matched control parasites. Multiple-testing correction was performed using the Benjamini–Hochberg method.

Genes were considered significantly differentially expressed when the false-discovery rate was below 0.05 and the absolute fold change exceeded 1.1. For strongly regulated genes, an additional absolute fold-change threshold of 1.5 was applied. Where fold change was expressed as lactate treatment relative to control, positive values consistently indicated increased expression following lactate exposure and negative values indicated reduced expression.

### Chromatin immunoprecipitation and quantitative PCR (ChIP-qPCR)

Trophozoite NF54 parasites were cultured as for Cut&Tag, with three independent biological replicates for each condition. The procedure followed [4], with H3K18La antibody used as the sole immunoprecipitating antibody.

Quantitative PCR was performed using gene-specific primers in technical duplicate for each biological replicate on a QuantStudio 6 Pro system using SensiFAST™ SYBR® Lo-ROX Kit (Meridian Bioscience, BIO-94005). Reactions contained approximately 3–4 ng immunoprecipitated or input DNA. Cycling conditions comprised an initial incubation for 2 min at 95°C followed by 40 cycles of 5 s at 95°C and 30 s at 60°C. Melt-curve analysis was used to assess amplification specificity.

Enrichment was calculated using the percentage-input method:

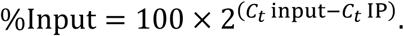

Technical replicate values were averaged to generate one measurement per biological replicate. For the gene-group comparison, replicate measurements were summarized to obtain a mean value for each gene, then grouped into the predefined control and target gene sets..

### Gene Ontology enrichment analysis

Gene Ontology enrichment analyses were performed using R (v4.5.1) for gene sets derived from RNA-seq, CUT&Tag and the integrated chromatin–transcription analyses.

Gene-to-GO annotations were obtained from the *P. falciparum* genome annotation and PlasmoDB resources [36]. Enrichment was assessed separately for Biological Process, Cellular Component and Molecular Function ontologies. The background universe was selected to match the analysis being performed. Genome-wide or peak-association analyses used the corresponding annotated gene universe, RNA-seq analyses used genes retained after expression filtering, and stage-comparison analyses used genes for which valid signal measurements were available in the relevant conditions. This avoided testing against genes that could not have entered the corresponding foreground set.

GO enrichment was evaluated using over-representation testing. For each term, the number of foreground genes annotated to that term was compared with the corresponding frequency in the selected background universe. Terms with nominal enrichment *P* values at or below 0.05 were retained for downstream summarization unless a more stringent criterion was specified in the individual analysis. Where adjusted values were generated, Benjamini–Hochberg correction was applied. Gene ratio was calculated as the number of genes from the analysed foreground set assigned to a GO term divided by the total number of genes in that foreground set.

Redundant GO terms were summarized using REVIGO rrvgo package (v. 1.16.0) REVIGO RREVIGO output fields describing term uniqueness and dispensability were scaled and used to calculate pairwise distances between terms. Classical multidimensional scaling was then used to project the terms into two-dimensional semantic space. The semantic-space coordinates therefore represent relative functional similarity and do not have a direct biological unit. Terms appearing close together are more semantically related than terms positioned farther apart.

For the REVIGO plots, representative terms were selected using the exported REVIGO Value ranking field according to the number retained in each final panel. Term labels were wrapped and positioned using label-repulsion algorithms solely to improve readability; label positioning did not affect the semantic coordinates.

For field-isolate comparisons, genes associated with H3K18La peaks were first identified separately in the six isolates. Shared sets were defined as genes present in at least four, five or six of all six isolates.

### Statistical analysis

Statistical analyses and graphical summaries were generated in R. Biological replicates, rather than technical replicates (in the case of ChIP-qPCR), were treated as independent observations. Technical replicates were averaged before statistical testing. The number of biological replicates used for each experiment is reported in the relevant Methods section and figure legend.

Two-group comparisons of gene-level or region-level measurements were performed using two-sided Wilcoxon rank-sum tests unless otherwise specified. Comparisons of control and 25 mM lactate conditions within *var*-gene subclasses were performed separately for each histone mark and subclass, followed by Benjamini–Hochberg correction across the related comparisons.

Associations between continuous variables were assessed using Spearman’s rank-correlation coefficient; the reported correlation coefficients and *P* values were derived from the Spearman tests rather than from the fitted linear models.

Principal component analysis was performed after log₂(signal + 1) transformation of gene-level promoter values. The matrix was centred and scaled before decomposition. Genes or samples with incomplete measurements were removed from the corresponding analysis.

## List of primers

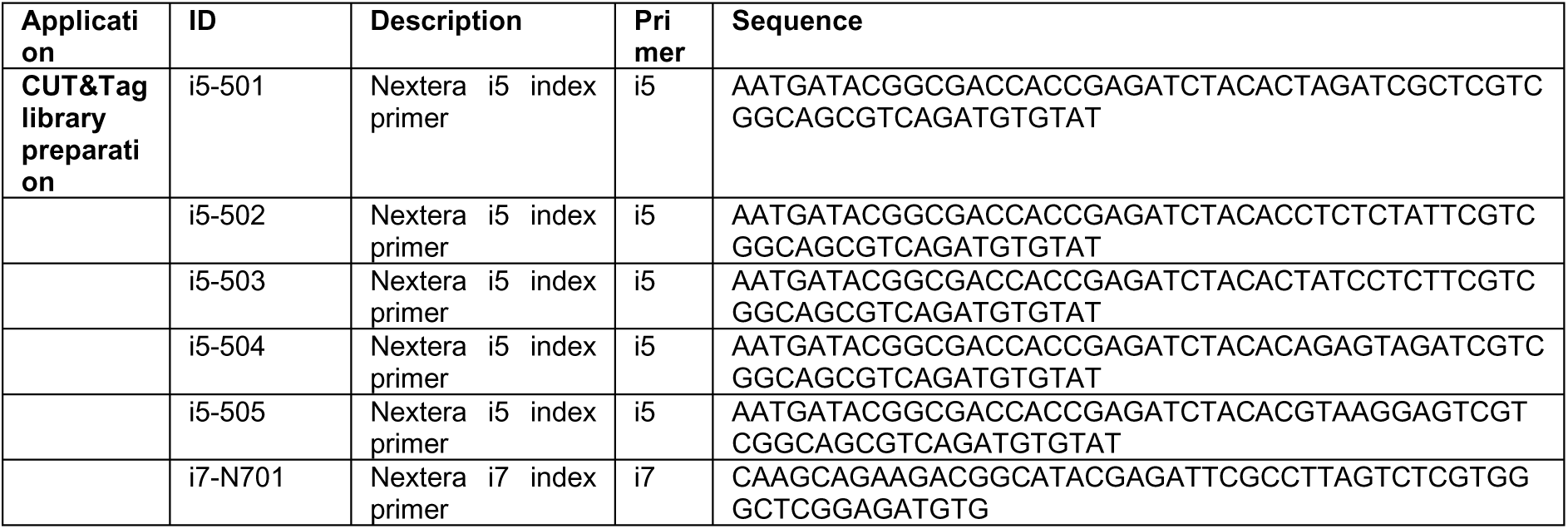
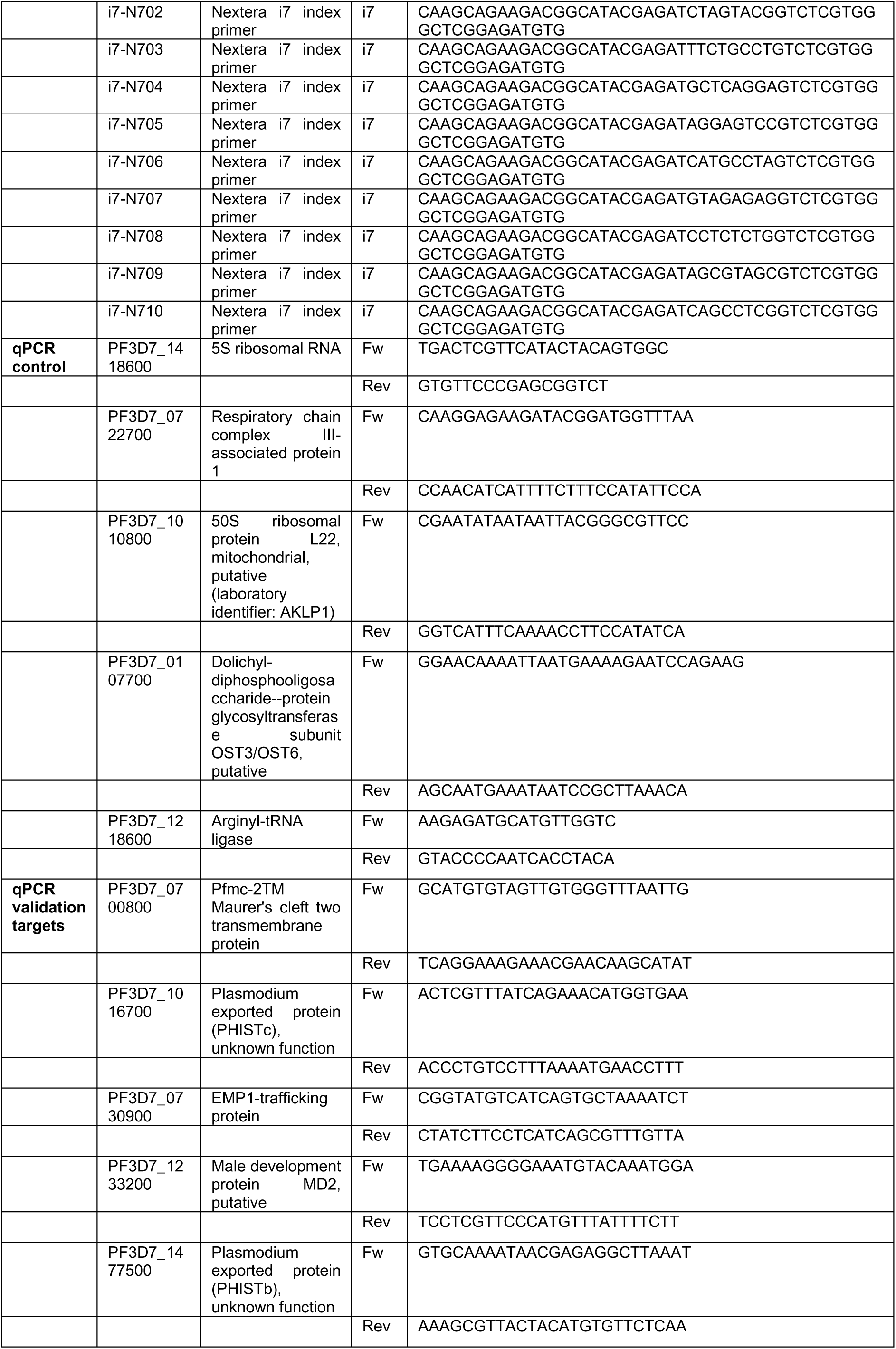
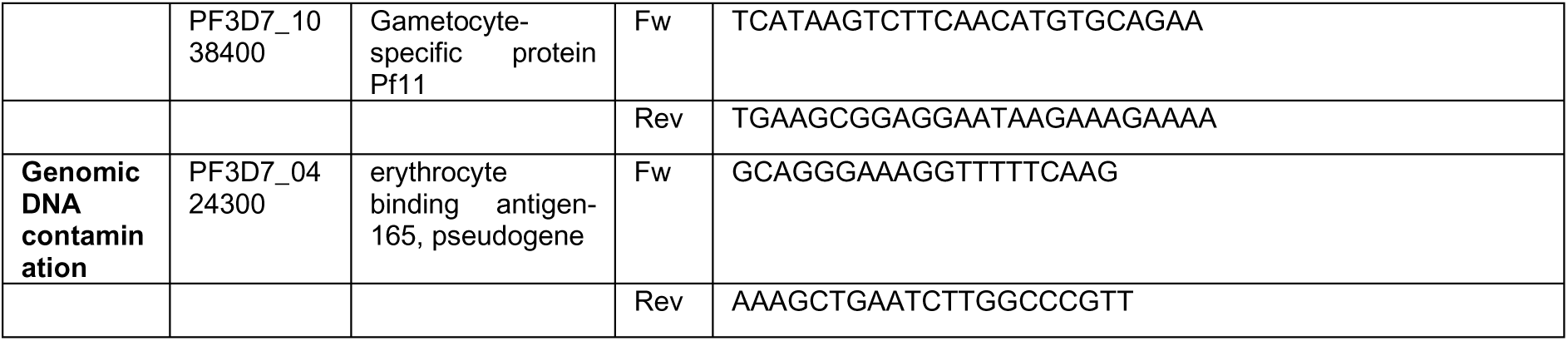

## List of antibodies

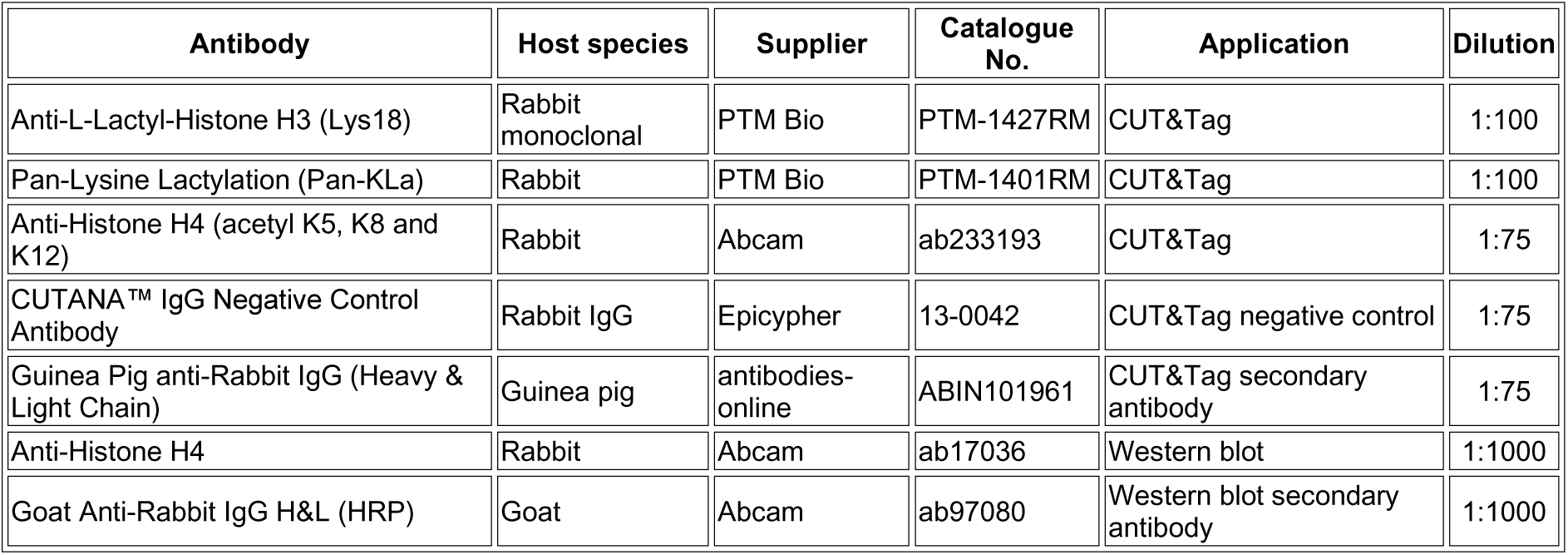

## Acknowledgements

We are grateful to Richard Bartfai & Jonas Gockel for advice on the Cut&Tag protocol, to Cambridge Genomic Services for help with sequencing, and to all study participants in The Gambia for consenting to participate in this study. We acknowledge the clinical staff at Brikama Health Centre and the laboratory staff at MRCG at LSHTM for their support with participant recruitment, sample collection and processing, particularly Fatoumata Bojang and Simon Correa.

This work was funded by Wellcome Discovery Award 225171/Z/22/Z to CJM, Royal Society International Exchange award IES\R2\252273 to CJM and AAN, and the UKRI Research England International Science Partnerships Fund (ISPF) Institutional Support Grant for Official Development Assistance to Cambridge University. Work at MRCG at LSHTM was also supported by the H3Africa Pan-African Malaria Genetic Epidemiology Network (PAMGEN), funded by the Science for Africa Foundation (grant H3A/18/002) to AAN. HM was supported by a WACCBIP DELTAS II Postdoctoral Fellowship (WACCBIP+NCDs: Awandare). The funders had no role in study design, data collection, interpretation or the decision to submit the work for publication.

