## Supplementary material for "Lactylated histones mark virulence gene families in the malaria parasite *Plasmodium falciparum*": Fig S

#### Supplementary Figure Legends

##### Figure S1. Quality control of trophozoite CUT&Tag datasets

**(A)** Representative immunoblot of H3K18La in control parasites and parasites exposed to 25 mM lactate for 12 h, with total H4 used as the loading control. The accompanying quantification shows H3K18La signal intensity relative to H4: bars represent the mean of three biological replicates, error bars indicate SEM and individual replicates are shown as points.

**(B)** Principal component analysis of genome-wide CPM-normalised CUT&Tag signal for H3K18La, H4K12La, pan-lysine lactylation (KLa) and H4Ac. PCA was performed independently for each histone mark using the corresponding deepTools coordinate output. Colours distinguish control and 25 mM lactate-treated parasites, and point shapes identify biological replicates.

**(C)** Distributions of SEACR peak widths for each histone mark, condition and biological replicate. Violin plots show the peak-width distributions and embedded boxplots indicate the median and interquartile range. Peak width is displayed on a log10 scale.

**(D)** Numbers of SEACR peaks detected for each histone mark under control and 25 mM lactate conditions. Bars represent the mean across three biological replicates, error bars indicate SEM and individual replicates are shown as points.

**(E)** Numbers of unique peak-associated genes detected for each histone mark and condition. A gene was counted once per sample after exclusion of assignments restricted to coding sequence or 3'UTR. Bars represent the mean across biological replicates, error bars indicate SEM and individual replicates are shown as points.

**(F)** Fraction of reads in peaks (FRiP) for each histone mark and condition. Bars represent the mean across biological replicates, error bars indicate SEM and individual replicates are shown as points.

**(G)** IgG negative-control CUT&Tag profiles across representative chromosomes 1, 7 and 14 in control and 25 mM lactate-treated parasites. Each replicate track was capped at its 99.5th percentile and scaled to its own dynamic range before replicate signals were averaged by condition. The tracks therefore illustrate the genomic distribution of background signal and are not intended for comparison of absolute signal intensity between conditions.

**(H)** GC-content versus local fragment-depth profiles for all trophozoite CUT&Tag libraries. The 14 nuclear chromosomes of the *P. falciparum* 3D7 reference genome were divided into complete, non-overlapping 300-bp windows; terminal partial windows were discarded. GC fraction and the number of overlapping properly paired primary fragments were calculated for each window, and windows were grouped into GC-fraction bins of width 0.01. Boxplots show the distribution of fragment counts within each GC bin. Outlier points are not displayed, and each library is shown using an independently adjusted linear y-axis to accommodate differences in sequencing depth.

**Figure S2. Gene Ontology programmes associated with CUT&Tag peaks and lactate-responsive chromatin changes**

**(A; first page)** REVIGO semantic-space representation of Gene Ontology Biological Process and Molecular Function terms enriched among all peak-associated genes for H3K18La, H4K12La, KLa and H4Ac. The ten most significant non-redundant terms are displayed for each histone mark and ontology.

**(B; second page)** REVIGO semantic-space representation of Biological Process, Molecular Function and Cellular Component terms enriched among genes associated with CUT&Tag regions showing increased signal following exposure to 25 mM lactate. Analyses are shown independently for H3K18La, H4K12La, KLa and H4Ac.

**(C; third page)** REVIGO semantic-space representation of Biological Process, Molecular Function and Cellular Component terms enriched among genes associated with CUT&Tag regions showing decreased signal following lactate treatment, shown independently for each histone mark. Panels labelled "No enriched GO terms" indicate comparisons in which no terms passed the applied enrichment criteria ($P.adj < 0.05$); panels containing a single term indicate that only one enriched term was available and multidimensional semantic reduction was therefore not applicable.

For all panels, redundant GO terms were reduced using REVIGO with the SIMREL semantic-similarity measure and a redundancy cutoff of 0.5. The ten highest-ranking terms were selected according to enrichment significance. Coordinates were generated by classical multidimensional scaling of distances calculated from scaled REVIGO Uniqueness and Dispensability values; spatial proximity therefore indicates semantic relatedness and does not represent effect size. Bubble size represents REVIGO GO-term size (LogSize), bubble-outline thickness denotes relative significance among the terms displayed in the corresponding panel, and bubble colour identifies the assigned biological category.

**Figure S3. Quality control of the trophozoite RNA-seq dataset**

**(A)** PCR-based confirmation of the absence of genomic DNA contamination in the cDNA preparations used for RNA sequencing. Primers targeting the intron-containing gene **PFD115w** were designed to span an intron, thereby generating products of different sizes from genomic DNA and spliced cDNA. Each biological replicate was analysed in two technical PCR replicates, shown as repeated R1, R2 and R3 lanes for control and 25 mM lactate-treated samples. All RNA-derived cDNA samples produced the smaller product expected following intron removal, whereas the genomic DNA control produced the larger intron-containing product. The absence of the larger genomic DNA-sized product from the cDNA lanes indicates that detectable genomic DNA contamination was not present. A DNA ladder is shown on the left, with the 75- and 200-bp positions indicated.

**(B)** Principal component analysis of $\log_2$(TPM + 1)-transformed gene-expression values. Genes with zero variance were removed, and the remaining variables were centred and scaled before PCA. Colours distinguish control and 25 mM lactate-treated samples, point shapes identify biological replicates, and axis labels indicate the percentage of total variance explained.

**(C)** RNA-seq library sizes, calculated as the total number of gene-assigned reads in each biological replicate.

**(D)** Numbers of genes before and after expression filtering. Genes with a summed raw read count greater than 10 across the six libraries were retained for downstream analysis. Values above the bars indicate the corresponding numbers of genes.

**(E)** Distributions of raw read counts and TPM values across biological replicates after $\log_2$(value + 1) transformation. Boxplots show the median and interquartile range; outlier points are not displayed.

**(F)** Distribution of the number of annotated transcripts per gene derived from the transcript-to-gene mapping table. Transcript–gene pairs were deduplicated before counting. Values above the bars indicate the number of genes with one, two or three annotated transcripts.

**Figure S4. Gene Ontology enrichment associated with genes showing concordant changes in RNA-seq/CUT&Tag responses, and ChIP-qPCR validation**

**(A)** REVIGO semantic-space representation of Cellular Component GO terms enriched among genes showing concordant RNA-seq and CUT&Tag changes following lactate treatment. The upper row shows genes with concordant increases and the lower row shows genes with concordant decreases for H3K18La, H4K12La, KLa and H4Ac. Up to ten highest-ranking non-redundant terms are displayed for each comparison. Spatial proximity indicates semantic similarity, bubble size represents GO-term size, bubble-outline thickness represents relative significance among displayed terms and bubble colour identifies the assigned biological category. The H4Ac concordant-down panel indicates that no Cellular Component terms passed the enrichment criteria.

**(B)** ChIP-qPCR validation using the H3K18La antibody. Left panel, relative H3K18La enrichment for control genes (without concordant RNA-seq/CUT&Tag changes) compared with selected target genes showing concordant lactate-associated changes. Bars represent the mean of the gene-level means, error bars indicate SEM and points show the individual gene means. Statistical significance was assessed using a two-sided Wilcoxon rank-sum test on gene-level means ($P = 0.00433$). Right panel, relative enrichment for each control and target gene. Bars show the mean across the available qPCR measurements, error bars indicate SEM and points represent individual measurements.

Control genes were **5SrRNA** (PF3D7_1418600; 5S ribosomal RNA), **AKLP1** (PF3D7_1010800; 50S ribosomal protein L22, mitochondrial, putative; laboratory identifier AKLP1), **Argnyl** (PF3D7_1218600; arginyl-tRNA ligase), **C3AP1** (PF3D7_0722700; respiratory chain complex III-associated protein 1), and **OST** (PF3D7_0107700; dolichyl-diphosphooligosaccharide–protein glycosyltransferase subunit OST3/OST6, putative). CUT&Tag up-regulated genes were **pfmc-2TM** (PF3D7_0700800; Pfmc-2TM Maurer's cleft two-transmembrane protein), **PHISTc** (PF3D7_1016700; *Plasmodium* exported protein, PHISTc), **EMP1** (PF3D7_0730900; EMP1-trafficking protein), **MD2** (PF3D7_1233200; male development protein MD2, putative), **PHISTb** (PF3D7_1477500; *Plasmodium* exported protein, PHISTb), and **Pf11** (PF3D7_1038400; gametocyte-specific protein Pf11).

**Figure S5. Experimental workflow and quality control of field-isolate and NF54 ring-stage CUT&Tag datasets**

**(A)** Schematic of the workflow used to process *P. falciparum*-infected clinical blood samples, comprising washing, dilution, leukocyte depletion, parasite release, extensive washing, fixation, cryopreservation, nuclei isolation, H3K18La CUT&Tag and sequencing.

**(B)** Genome-wide PCA of sample-level BigWig signal quantified in equal fixed genomic bins. The left plot compares H3K18La profiles from the six field isolates with control and 25 mM lactate-treated NF54 ring-stage parasites; the right plot compares KLa profiles from control and lactate-treated NF54 ring-stage parasites. PCA was performed independently for the two comparisons after removal of invariant bins, centring and scaling. Field isolates are shown individually, whereas colours and point shapes distinguish NF54 treatment conditions and biological replicates.

**(C)** Distributions of SEACR peak widths in the six field-isolate H3K18La datasets and in NF54 ring-stage H3K18La and KLa datasets. Violin plots show the peak-width distributions and embedded boxplots indicate the median and interquartile range. Peak width is displayed on a $\log_{10}$ scale.

**(D)** Numbers of SEACR peaks detected in each field isolate and in NF54 ring-stage H3K18La and KLa datasets. Field isolates are shown individually. For NF54 samples, bars represent the mean across three biological replicates, error bars indicate SEM and points show individual replicates.

**(E)** Numbers of unique peak-associated genes. A gene was counted once per sample after exclusion of assignments restricted to coding sequence or 3'UTR. Field isolates are shown individually; NF54 bars represent mean ± SEM across three biological replicates with individual replicates overlaid.

**(F)** FRiP scores calculated from sample-matched BAM files and SEACR peaks. Properly paired, primary mapped fragments were counted once using first mates; unmapped, duplicate, secondary, supplementary and QC-failed alignments were excluded. FRiP was calculated as the number of filtered fragments overlapping at least one relaxed peak divided by the total number of filtered fragments. Field isolates are shown individually and NF54 bars represent mean ± SEM with biological replicates overlaid.

**(G; continuation page)** GC-content versus local fragment-depth profiles for NF54 ring-stage H3K18La and KLa libraries and the six field-isolate H3K18La datasets. Complete non-overlapping 300-bp windows from the 14 nuclear chromosomes were assigned to 0.01-wide GC-fraction bins, and overlapping properly paired primary fragments were counted for each window. Boxplots show fragment-count distributions within each GC bin. Outlier points are not displayed, and each library is shown using an independently adjusted linear y-axis.

**Figure S6. Functional programmes shared among field isolates and promoter H3K18La distributions across var subclasses**

**(A)** Gene Ontology enrichment across genes detected in increasing numbers of field-isolate datasets. Columns indicate genes shared by one to six of the six field isolates. Biological Process, Cellular Component and Molecular Function terms were analysed independently. For each shared-set and ontology combination, the five most significant terms were selected; the union of selected terms is displayed within each ontology, and a dot is shown only when the corresponding term was selected for that shared-set comparison. Dot fill represents −log10(P), with darker shading indicating stronger enrichment; dot size is constant.

**(B)** Distribution of H3K18La signal upstream of the TSS across non-*var* genes and *var* genes belonging to the UPSA, UPSB, UPSC and UPSE promoter classes. The analysis includes the six field isolates and NF54 ring- and trophozoite-stage parasites under control and 25 mM lactate conditions. Only rows assigned to the promoter region were retained. Signals are expressed as gene-level log2(CUT&Tag signal + 1). Violin plots show the gene distributions and embedded boxplots indicate the median and interquartile range. A common y-axis scale is used across all samples.

**Figure S7. Stage-specific promoter lactylation and functional enrichment of ring-stage peak-associated genes**

**(A)** Gene-level comparison of promoter-associated H3K18La and KLa CUT&Tag signal between NF54 ring and trophozoite stages under control and 25 mM lactate conditions. Each point represents one gene and signals are shown as log2(signal + 1). The red dashed line indicates equality between ring- and trophozoite-stage signal, and the blue line represents a fitted linear regression for visualisation. Spearman correlation coefficients and associated P values are shown within each panel.

**(B)** REVIGO semantic-space representation of Biological Process and Molecular Function terms enriched among all ring-stage genes associated with H3K18La or KLa peaks. The eight most significant non-redundant terms are displayed for each histone mark and ontology.

**(C)** REVIGO semantic-space representation of Biological Process, Molecular Function and Cellular Component terms enriched among ring-stage genes associated with lactate-induced H3K18La or KLa CUT&Tag signal. The eight most significant non-redundant terms are displayed for each histone mark and ontology.

For panels B and C, coordinates were generated by classical multidimensional scaling of distances calculated from scaled REVIGO Uniqueness and Dispensability values. Spatial proximity indicates semantic similarity and does not represent enrichment magnitude. Bubble size represents GO-term size, bubble-outline thickness denotes relative significance among displayed terms and bubble colour identifies the assigned biological category.

#### Supplementary Table Legends

**Supplementary Data 1. CUT&Tag peak and differential enrichment analyses in NF54 trophozoites.** Condition-specific SEACR-called peaks and DESeq2 differential enrichment analyses for H3K18La, H4K12La, KLa and H4Ac in control and 25 mM lactate-treated samples.

**Supplementary Data 2. Gene Ontology enrichment of histone modification-associated genes in NF54 trophozoites.** GO enrichment results and REVIGO-reduced functional annotations for genes associated with H3K18La, H4K12La, KLa and H4Ac.

**Supplementary Data 3. Transcriptomic response of NF54 trophozoites to lactate treatment.** Gene-level RNA-seq counts, TPM values and differential expression statistics, together with GO enrichment and REVIGO analyses of up- and down-regulated genes following 25 mM lactate treatment.

**Supplementary Data 4. Concordance between histone modifications and transcriptional responses to lactate.** Gene-level concordance between changes in H3K18La, H4K12La, KLa or H4Ac enrichment and RNA expression following lactate treatment, together with GO enrichment of concordantly regulated gene sets.

**Supplementary Data 5. CUT&Tag peak and differential enrichment analyses in NF54 rings**. H3K18La and KLa peak annotations and differential peak enrichment analyses, together with GO and REVIGO analyses of condition-specific, merged and lactate-induced peak-associated gene sets.

**Supplementary Data 6. CUT&Tag peak conservation across *P. falciparum* field isolates.** Annotated CUT&Tag peaks from six *P. falciparum* field isolates and genomic peak sets stratified according to their degree of sharing across one to six isolates, together with GO analyses of each stratified peak set.

**Fig. S1**

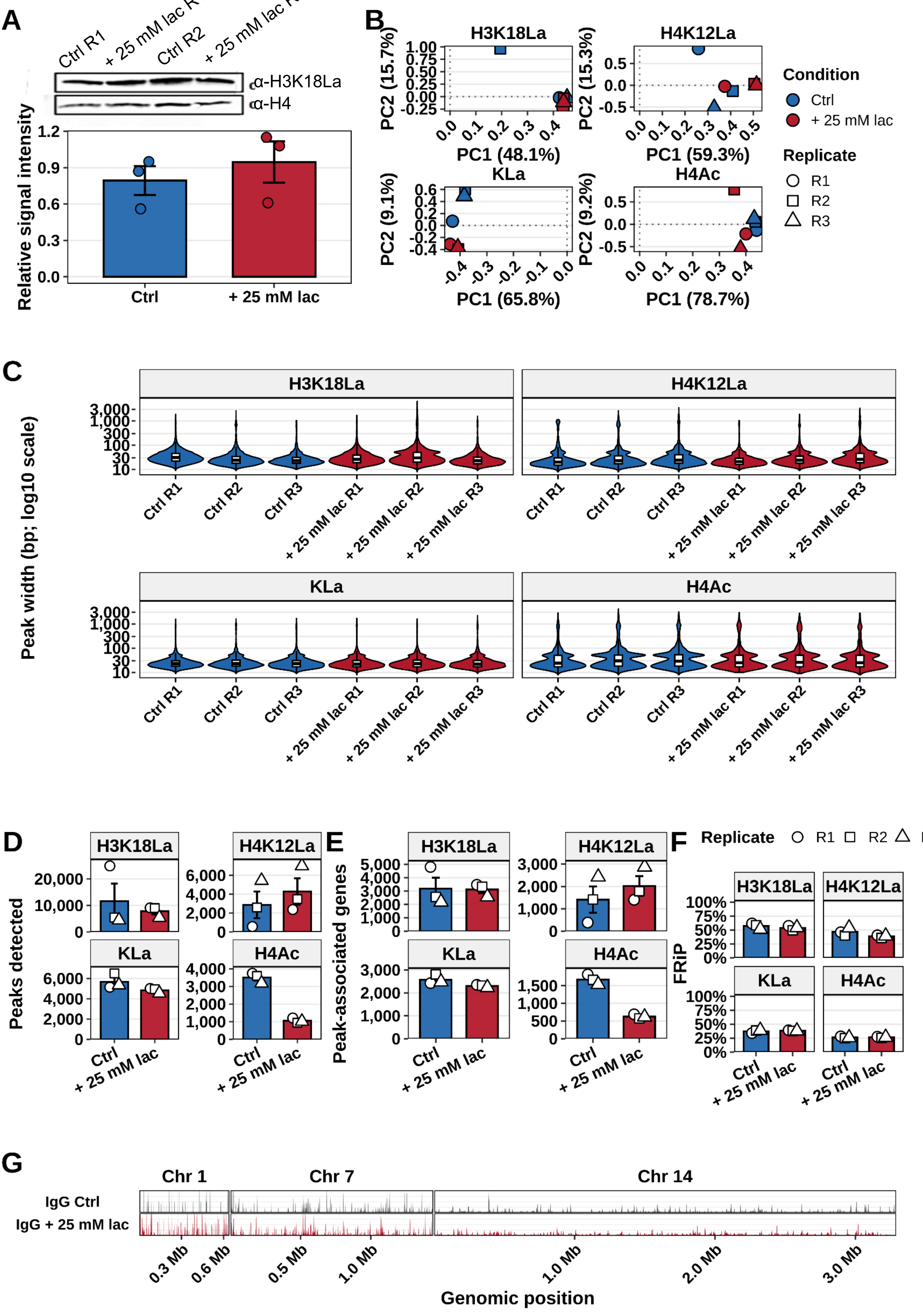
A
Ctrl R1
+ 25 mM lac R1
Ctrl R2
+ 25 mM lac R2
α-H3K18La
α-H4
Relative signal intensity
1.2
0.9
0.6
0.3
0.0
Ctrl
+ 25 mM lac
B
H3K18La
H4K12La
KLa
H4Ac
PC2 (15.7%)
PC2 (15.3%)
PC2 (9.1%)
PC2 (9.2%)
PC1 (48.1%)
PC1 (59.3%)
PC1 (65.8%)
PC1 (78.7%)
Condition
Ctrl
+ 25 mM lac
Replicate
R1
R2
R3
C
Peak width (bp; log10 scale)
3,000
1,000
300
100
30
10
Ctrl R1
Ctrl R2
Ctrl R3
+ 25 mM lac R1
+ 25 mM lac R2
+ 25 mM lac R3
D
Peaks detected
E
Peak-associated genes
F
FRiP
Replicate
R1
R2
G
Chr 1
Chr 7
Chr 14
IgG Ctrl
IgG + 25 mM lac
0.3 Mb
0.6 Mb
0.5 Mb
1.0 Mb
1.0 Mb
2.0 Mb
3.0 Mb
Genomic position

Fig. S1

H

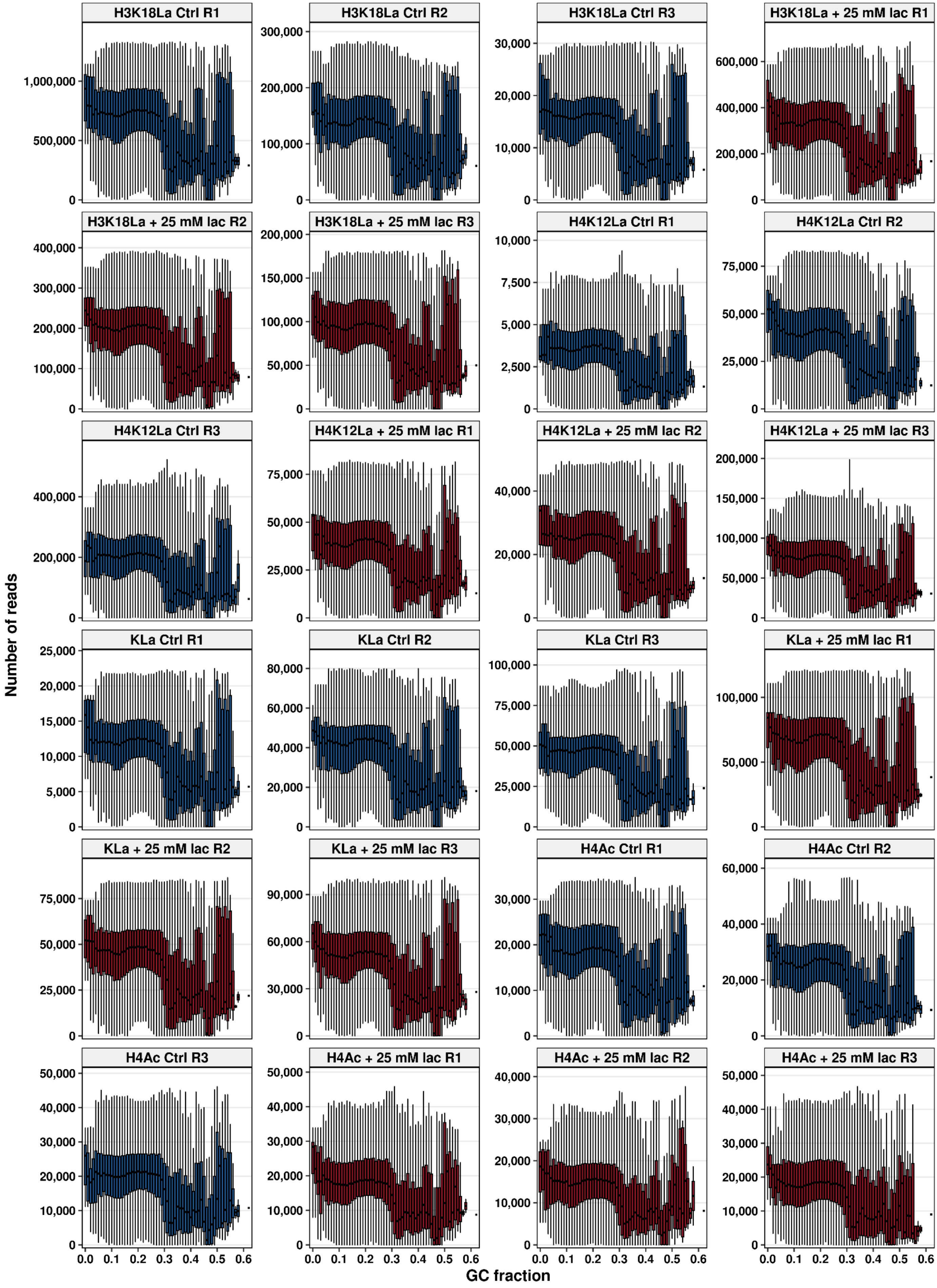

H3K18La Ctrl R1
H3K18La Ctrl R2
H3K18La Ctrl R3
H3K18La + 25 mM lac R1
H3K18La + 25 mM lac R2
H3K18La + 25 mM lac R3
H4K12La Ctrl R1
H4K12La Ctrl R2
H4K12La Ctrl R3
H4K12La + 25 mM lac R1
H4K12La + 25 mM lac R2
H4K12La + 25 mM lac R3
KLa Ctrl R1
KLa Ctrl R2
KLa Ctrl R3
KLa + 25 mM lac R1
KLa + 25 mM lac R2
KLa + 25 mM lac R3
H4Ac Ctrl R1
H4Ac Ctrl R2
H4Ac Ctrl R3
H4Ac + 25 mM lac R1
H4Ac + 25 mM lac R2
H4Ac + 25 mM lac R3
Number of reads
GC fraction
0.0 0.1 0.2 0.3 0.4 0.5 0.6

**Fig. S2**

**A**

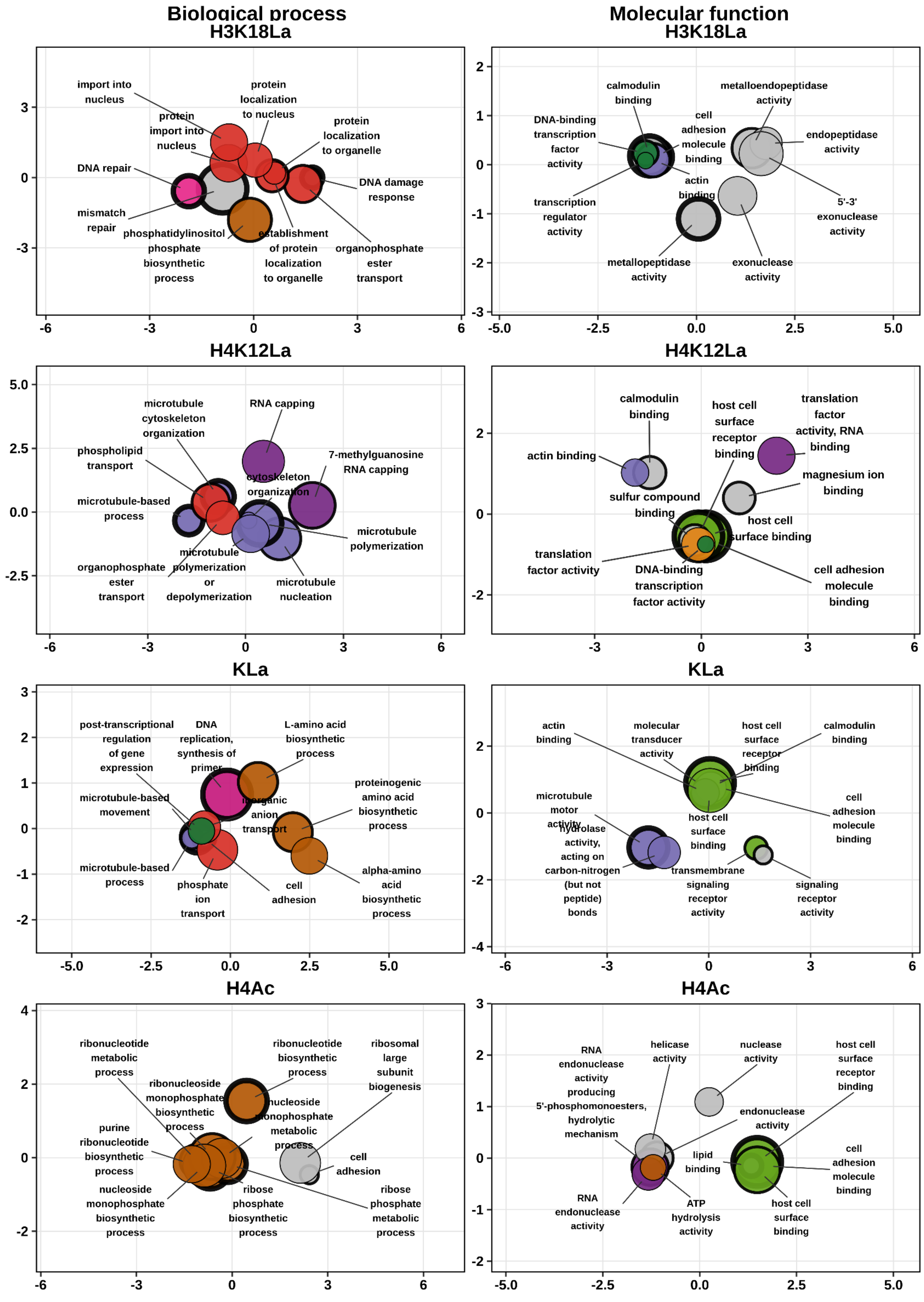
Biological process
Molecular function
H3K18La
import into nucleus
protein localization to nucleus
protein import into nucleus
protein localization to organelle
DNA repair
DNA damage response
mismatch repair
phosphatidylinositol phosphate biosynthetic process
establishment of protein localization to organelle
organophosphate ester transport
calmodulin binding
metalloendopeptidase activity
DNA-binding transcription factor activity
cell adhesion molecule binding
endopeptidase activity
actin binding
transcription regulator activity
5'-3' exonuclease activity
metallopeptidase activity
exonuclease activity
H4K12La
microtubule cytoskeleton organization
RNA capping
phospholipid transport
7-methylguanosine RNA capping
cytoskeleton organization
microtubule-based process
microtubule polymerization
organophosphate ester transport
microtubule polymerization or depolymerization
microtubule nucleation
calmodulin binding
host cell surface receptor binding
translation factor activity, RNA binding
actin binding
magnesium ion binding
sulfur compound binding
host cell surface binding
translation factor activity
DNA-binding transcription factor activity
cell adhesion molecule binding
KLa
post-transcriptional regulation of gene expression
DNA replication, synthesis of primer
L-amino acid biosynthetic process
proteinogenic amino acid biosynthetic process
microtubule-based movement
inorganic anion transport
microtubule-based process
phosphate ion transport
cell adhesion
alpha-amino acid biosynthetic process
actin binding
molecular transducer activity
host cell surface receptor binding
calmodulin binding
microtubule motor activity
cell adhesion molecule binding
hydrolase activity, acting on carbon-nitrogen (but not peptide) bonds
host cell surface binding
transmembrane signaling receptor activity
signaling receptor activity
H4Ac
ribonucleotide metabolic process
ribonucleotide biosynthetic process
ribosomal large subunit biogenesis
ribonucleoside monophosphate biosynthetic process
nucleoside monophosphate metabolic process
purine ribonucleotide biosynthetic process
cell adhesion
nucleoside monophosphate biosynthetic process
ribose phosphate biosynthetic process
ribose phosphate metabolic process
RNA endonuclease activity producing 5'-phosphomonoesters, hydrolytic mechanism
helicase activity
nuclease activity
host cell surface receptor binding
endonuclease activity
lipid binding
cell adhesion molecule binding
RNA endonuclease activity
ATP hydrolysis activity
host cell surface binding

Fig. S2

B

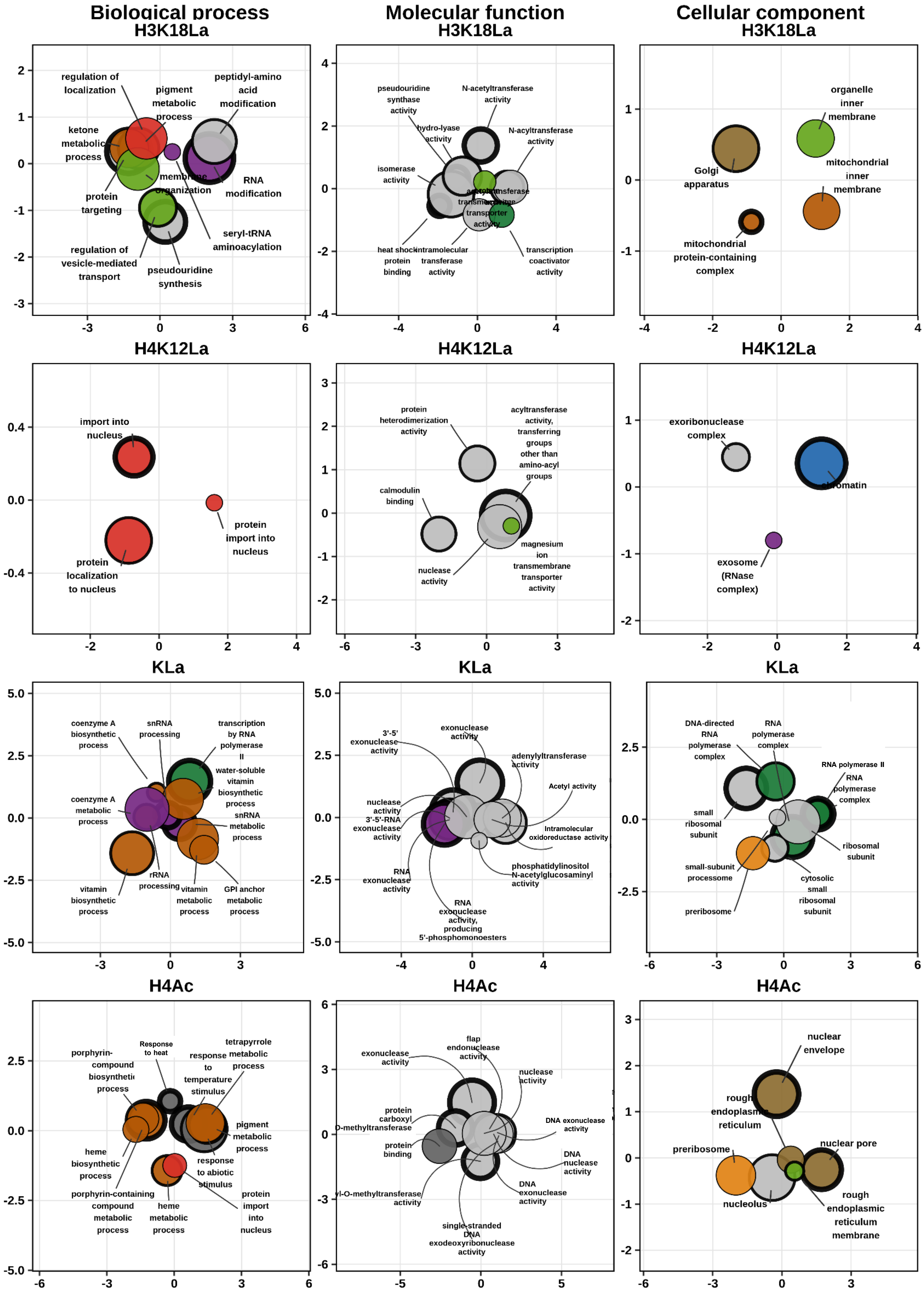

Biological process
Molecular function
Cellular component
H3K18La
regulation of localization
pigment metabolic process
peptidyl-amino acid modification
ketone metabolic process
membrane organization
RNA modification
protein targeting
seryl-tRNA aminoacylation
regulation of vesicle-mediated transport
pseudouridine synthesis
pseudouridine synthase activity
N-acetyltransferase activity
hydro-lyase activity
N-acyltransferase activity
isomerase activity
heat shock protein binding
intramolecular transferase activity
transcription coactivator activity
organelle inner membrane
Golgi apparatus
mitochondrial inner membrane
mitochondrial protein-containing complex
H4K12La
import into nucleus
protein import into nucleus
protein localization to nucleus
protein heterodimerization activity
acyltransferase activity, transferring groups other than amino-acyl groups
calmodulin binding
magnesium ion transmembrane transporter activity
nuclease activity
exoribonuclease complex
chromatin
exosome (RNase complex)
KLa
coenzyme A biosynthetic process
snRNA processing
transcription by RNA polymerase II
water-soluble vitamin biosynthetic process
coenzyme A metabolic process
snRNA metabolic process
vitamin biosynthetic process
rRNA processing
vitamin metabolic process
GPI anchor metabolic process
3'-5' exonuclease activity
exonuclease activity
adenylyltransferase activity
Acetyl activity
nuclease activity
3'-5'-RNA exonuclease activity
Intramolecular oxidoreductase activity
phosphatidylinositol N-acetylglucosaminyl activity
RNA exonuclease activity
RNA exonuclease activity, producing 5'-phosphomonoesters
DNA-directed RNA polymerase complex
RNA polymerase complex
RNA polymerase II
RNA polymerase complex
small ribosomal subunit
ribosomal subunit
small-subunit processome
cytosolic small ribosomal subunit
preribosome
H4Ac
porphyrin-compound biosynthetic process
Response to heat
response to temperature stimulus
tetrapyrrole metabolic process
pigment metabolic process
heme biosynthetic process
response to abiotic stimulus
porphyrin-containing compound metabolic process
heme metabolic process
protein import into nucleus
flap endonuclease activity
exonuclease activity
nuclease activity
protein carboxyl O-methyltransferase
DNA exonuclease activity
protein binding
nuclease activity
DNA exonuclease activity
single-stranded DNA exodeoxyribonuclease activity
nuclear envelope
rough endoplasmic reticulum
preribosome
nuclear pore
nucleolus
rough endoplasmic reticulum membrane

**Fig. S2**

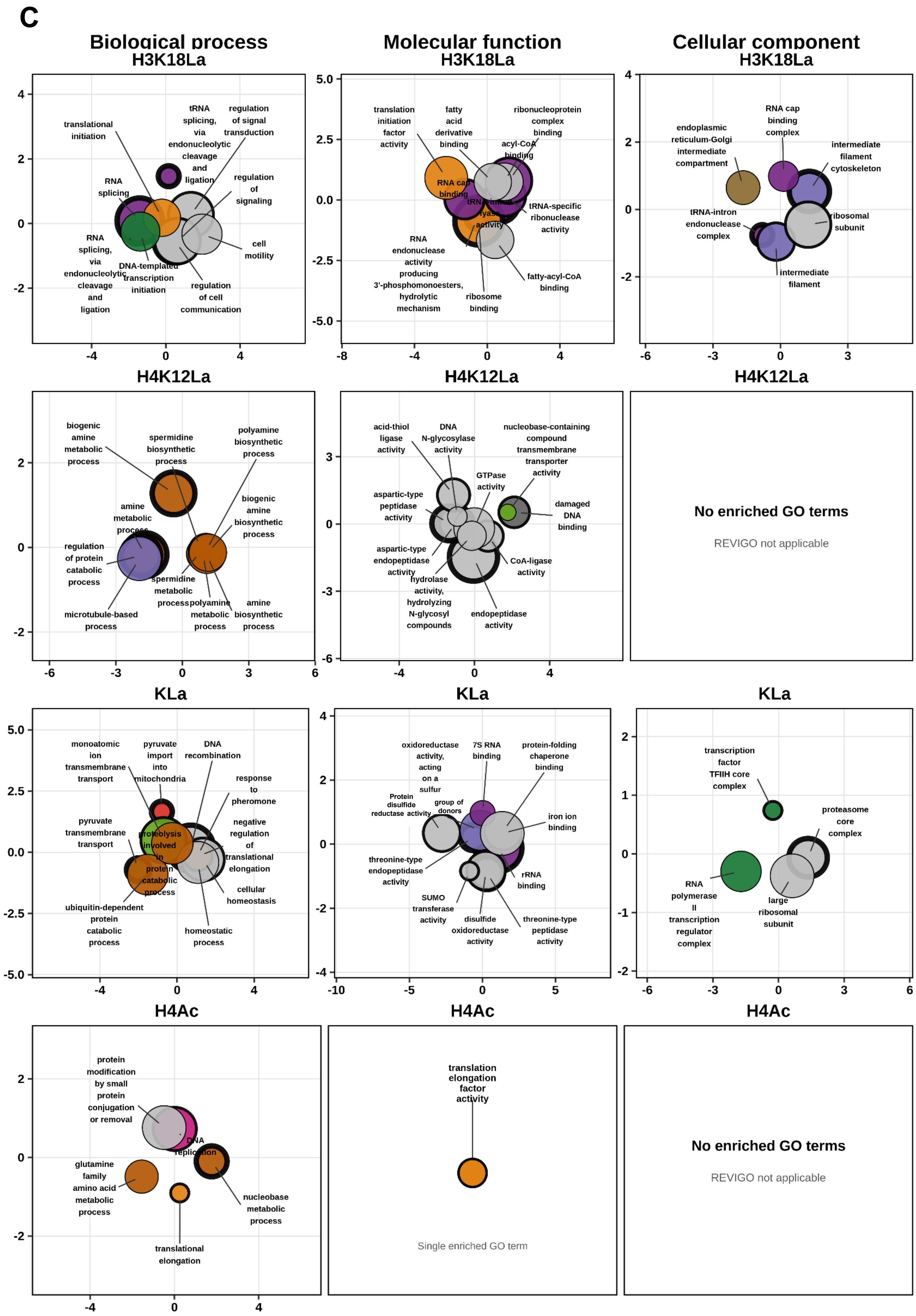

C
Biological process
H3K18La
translational initiation
tRNA splicing, via endonucleolytic cleavage and ligation
regulation of signal transduction
RNA splicing
regulation of signaling
RNA splicing, via endonucleolytic cleavage and ligation
DNA-templated transcription initiation
regulation of cell communication
cell motility
Molecular function
H3K18La
translation initiation factor activity
fatty acid derivative binding
ribonucleoprotein complex binding
acyl-CoA binding
RNA cap binding
tRNA-intron lyase activity
tRNA-specific ribonuclease activity
RNA endonuclease activity producing 3'-phosphomonoesters, hydrolytic mechanism
ribosome binding
fatty-acyl-CoA binding
Cellular component
H3K18La
endoplasmic reticulum-Golgi intermediate compartment
RNA cap binding complex
intermediate filament cytoskeleton
tRNA-intron endonuclease complex
ribosomal subunit
intermediate filament
H4K12La
biogenic amine metabolic process
spermidine biosynthetic process
polyamine biosynthetic process
amine metabolic process
biogenic amine biosynthetic process
regulation of protein catabolic process
spermidine metabolic process
polyamine metabolic process
amine biosynthetic process
microtubule-based process
H4K12La
acid-thiol ligase activity
DNA N-glycosylase activity
nucleobase-containing compound transmembrane transporter activity
GTPase activity
aspartic-type peptidase activity
damaged DNA binding
aspartic-type endopeptidase activity
CoA-ligase activity
hydrolase activity, hydrolyzing N-glycosyl compounds
endopeptidase activity
H4K12La
No enriched GO terms
REVIGO not applicable
KLa
monoatomic ion transmembrane transport
pyruvate import into mitochondria
DNA recombination
response to pheromone
pyruvate transmembrane transport
proteolysis involved in protein catabolic process
negative regulation of translational elongation
cellular homeostasis
ubiquitin-dependent protein catabolic process
homeostatic process
KLa
oxidoreductase activity, acting on a sulfur
7S RNA binding
protein-folding chaperone binding
Protein disulfide reductase activity
group of donors
iron ion binding
threonine-type endopeptidase activity
rRNA binding
SUMO transferase activity
disulfide oxidoreductase activity
threonine-type peptidase activity
KLa
transcription factor TFIIH core complex
proteasome core complex
RNA polymerase II transcription regulator complex
large ribosomal subunit
H4Ac
protein modification by small protein conjugation or removal
DNA replication
glutamine family amino acid metabolic process
nucleobase metabolic process
translational elongation
H4Ac
translation elongation factor activity
Single enriched GO term
H4Ac
No enriched GO terms
REVIGO not applicable

**Fig. S3**

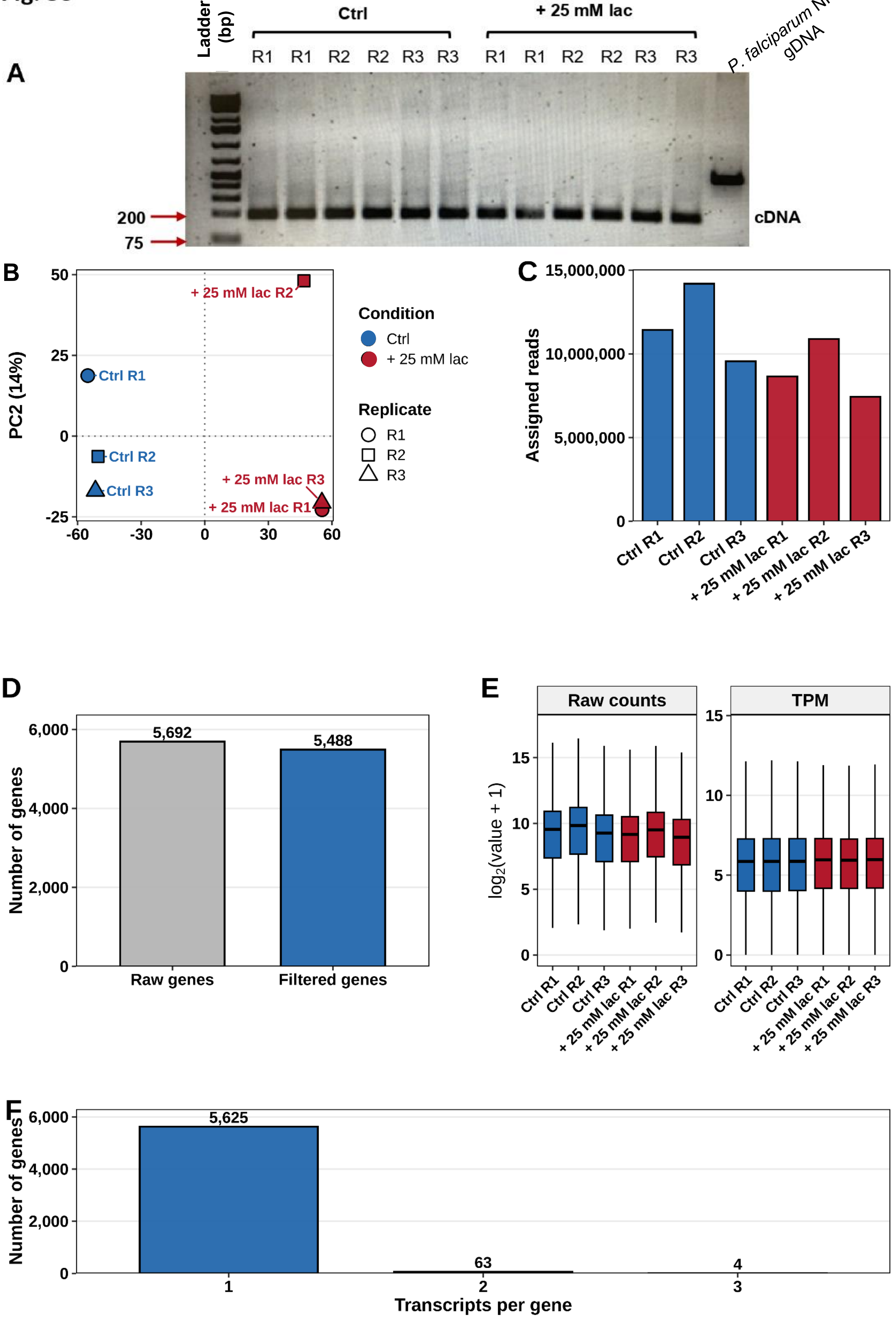
A
Ladder (bp)
Ctrl
+ 25 mM lac
R1 R1 R2 R2 R3 R3 R1 R1 R2 R2 R3 R3
P. falciparum NF54 gDNA
200
75
cDNA
B
PC2 (14%)
50
25
0
-25
-60 -30 0 30 60
+ 25 mM lac R2
Ctrl R1
Ctrl R2
Ctrl R3
+ 25 mM lac R3
+ 25 mM lac R1
Condition
Ctrl
+ 25 mM lac
Replicate
R1
R2
R3
C
Assigned reads
15,000,000
10,000,000
5,000,000
0
Ctrl R1
Ctrl R2
Ctrl R3
+ 25 mM lac R1
+ 25 mM lac R2
+ 25 mM lac R3
D
Number of genes
6,000
4,000
2,000
0
5,692
5,488
Raw genes
Filtered genes
E
Raw counts
TPM
log2(value + 1)
15
10
5
0
Ctrl R1
Ctrl R2
Ctrl R3
+ 25 mM lac R1
+ 25 mM lac R2
+ 25 mM lac R3
F
Number of genes
6,000
4,000
2,000
0
5,625
63
4
1
2
3
Transcripts per gene

Fig. S4

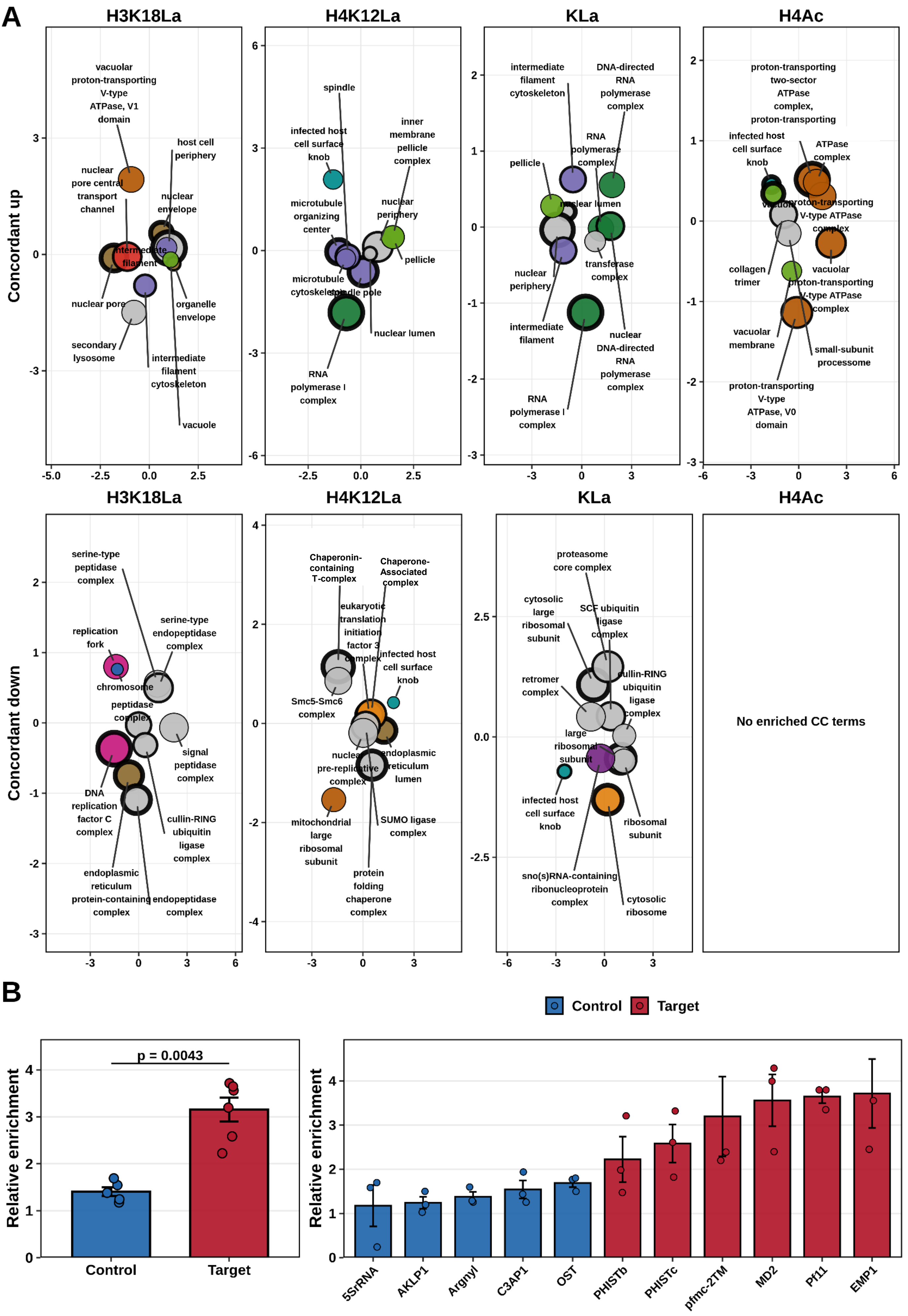
A
H3K18La
H4K12La
KLa
H4Ac
Concordant up
Concordant down
No enriched CC terms
B
Control
Target
Relative enrichment
p = 0.0043
5SrRNA
AKLP1
Argnyl
C3AP1
OST
PHISTb
PHISTc
pfmc-2TM
MD2
Pf11
EMP1

### Fig. S5

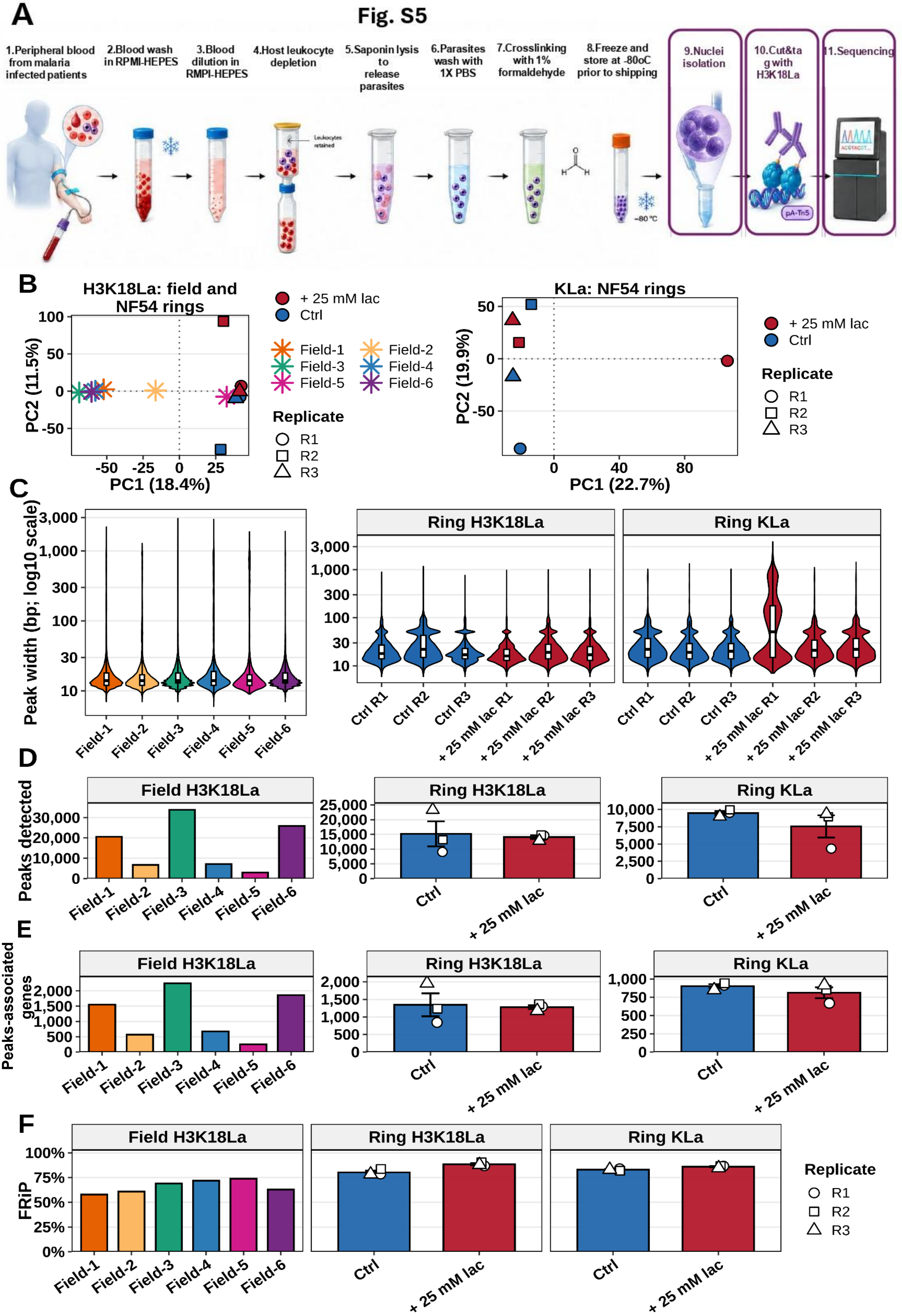

A
1.Peripheral blood from malaria infected patients
2.Blood wash in RPMI-HEPES
3.Blood dilution in RMPI-HEPES
4.Host leukocyte depletion
5.Saponin lysis to release parasites
6.Parasites wash with 1X PBS
7.Crosslinking with 1% formaldehyde
8.Freeze and store at -80oC prior to shipping
9.Nuclei isolation
10.Cut&tag with H3K18La
11.Sequencing
-80 °C
pA-Tn5
B
H3K18La: field and NF54 rings
PC1 (18.4%)
PC2 (11.5%)
+ 25 mM lac
Ctrl
Field-1
Field-2
Field-3
Field-4
Field-5
Field-6
Replicate
R1
R2
R3
KLa: NF54 rings
PC1 (22.7%)
PC2 (19.9%)
C
Peak width (bp; log10 scale)
Ring H3K18La
Ring KLa
Ctrl R1
Ctrl R2
Ctrl R3
+ 25 mM lac R1
+ 25 mM lac R2
+ 25 mM lac R3
D
Peaks detected
Field H3K18La
Ring H3K18La
Ring KLa
Ctrl
+ 25 mM lac
E
Peaks-associated genes
Field H3K18La
Ring H3K18La
Ring KLa
F
FRiP
Field H3K18La
Ring H3K18La
Ring KLa
Replicate
R1
R2
R3

Fig. S6

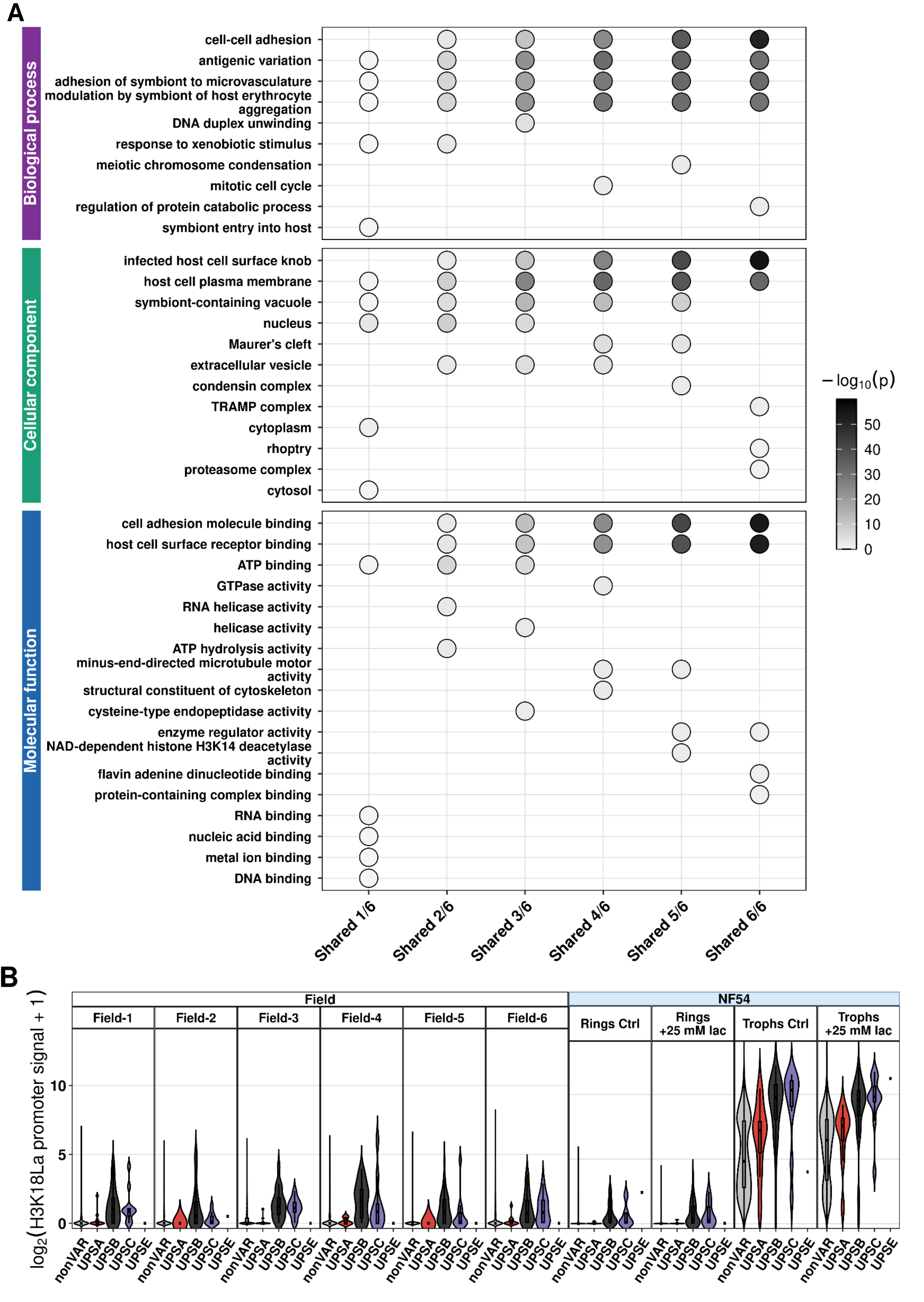

A
Biological process
cell-cell adhesion
antigenic variation
adhesion of symbiont to microvasculature
modulation by symbiont of host erythrocyte aggregation
DNA duplex unwinding
response to xenobiotic stimulus
meiotic chromosome condensation
mitotic cell cycle
regulation of protein catabolic process
symbiont entry into host
Cellular component
infected host cell surface knob
host cell plasma membrane
symbiont-containing vacuole
nucleus
Maurer's cleft
extracellular vesicle
condensin complex
TRAMP complex
cytoplasm
rhoptry
proteasome complex
cytosol
Molecular function
cell adhesion molecule binding
host cell surface receptor binding
ATP binding
GTPase activity
RNA helicase activity
helicase activity
ATP hydrolysis activity
minus-end-directed microtubule motor activity
structural constituent of cytoskeleton
cysteine-type endopeptidase activity
enzyme regulator activity
NAD-dependent histone H3K14 deacetylase activity
flavin adenine dinucleotide binding
protein-containing complex binding
RNA binding
nucleic acid binding
metal ion binding
DNA binding
Shared 1/6
Shared 2/6
Shared 3/6
Shared 4/6
Shared 5/6
Shared 6/6
$-\log_{10}(p)$
50
40
30
20
10
0
B
Field
NF54
Field-1
Field-2
Field-3
Field-4
Field-5
Field-6
Rings Ctrl
Rings +25 mM lac
Trophs Ctrl
Trophs +25 mM lac
$\log_2$(H3K18La promoter signal + 1)
10
5
0
nonVAR
UPSA
UPSB
UPSC
UPSE

Fig. S7

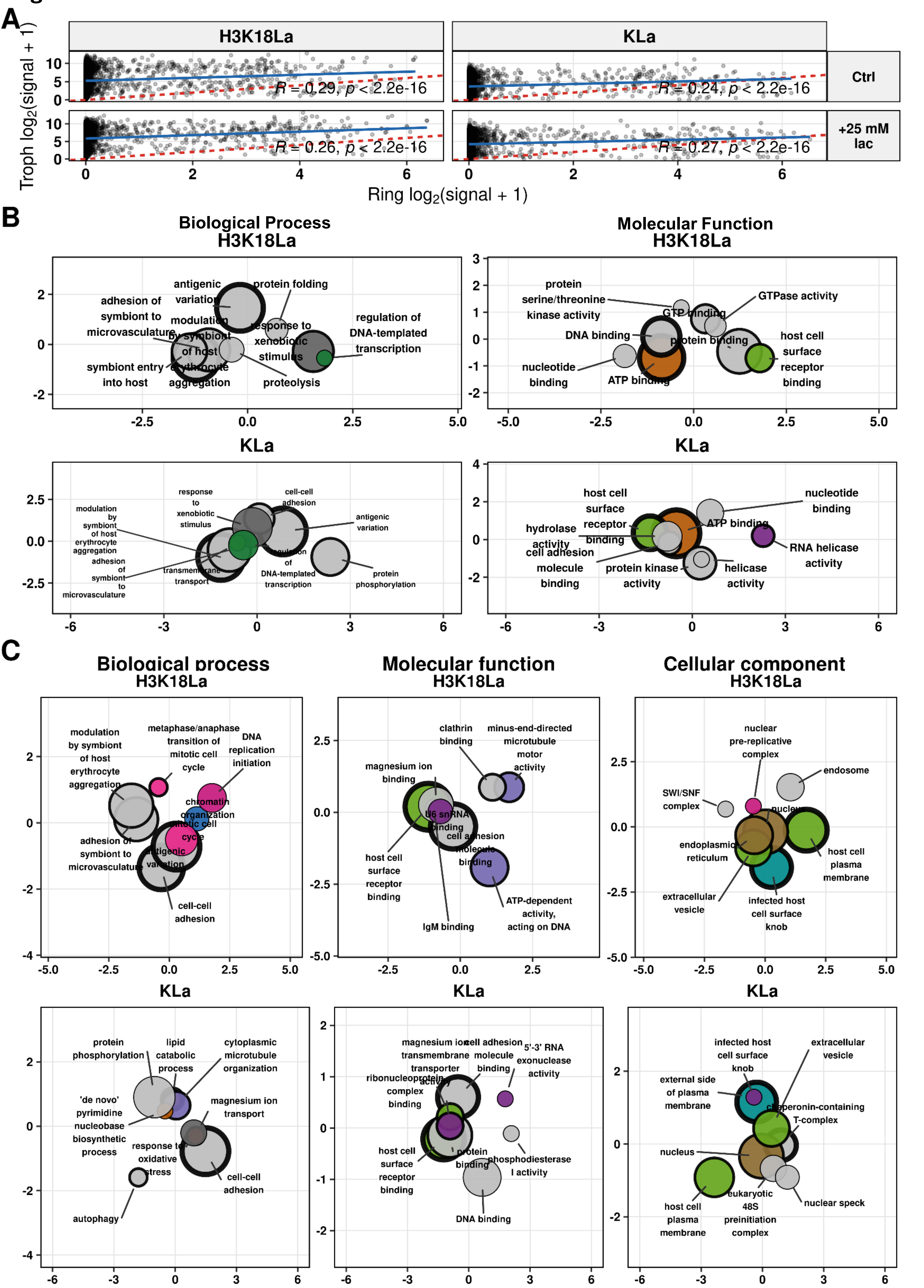
A
H3K18La
KLa
Ctrl
+25 mM lac
Troph log2(signal + 1)
Ring log2(signal + 1)
R = 0.29, p < 2.2e-16
R = 0.24, p < 2.2e-16
R = 0.26, p < 2.2e-16
R = 0.27, p < 2.2e-16
B
Biological Process
H3K18La
Molecular Function
H3K18La
antigenic variation
protein folding
adhesion of symbiont to microvasculature
modulation by symbiont of host erythrocyte aggregation
response to xenobiotic stimulus
regulation of DNA-templated transcription
symbiont entry into host
proteolysis
protein serine/threonine kinase activity
GTP binding
GTPase activity
DNA binding
protein binding
host cell surface receptor binding
nucleotide binding
ATP binding
KLa
response to xenobiotic stimulus
cell-cell adhesion
modulation by symbiont of host erythrocyte aggregation
antigenic variation
adhesion of symbiont to microvasculature
transmembrane transport
regulation of DNA-templated transcription
protein phosphorylation
host cell surface receptor binding
nucleotide binding
hydrolase activity
ATP binding
RNA helicase activity
cell adhesion molecule binding
protein kinase activity
helicase activity
C
Biological process
H3K18La
Molecular function
H3K18La
Cellular component
H3K18La
modulation by symbiont of host erythrocyte aggregation
metaphase/anaphase transition of mitotic cell cycle
DNA replication initiation
chromatin organization
mitotic cell cycle
adhesion of symbiont to microvasculature
antigenic variation
cell-cell adhesion
clathrin binding
minus-end-directed microtubule motor activity
magnesium ion binding
U6 snRNA binding
cell adhesion molecule binding
host cell surface receptor binding
ATP-dependent activity, acting on DNA
IgM binding
nuclear pre-replicative complex
endosome
SWI/SNF complex
nucleus
endoplasmic reticulum
host cell plasma membrane
extracellular vesicle
infected host cell surface knob
KLa
protein phosphorylation
lipid catabolic process
cytoplasmic microtubule organization
'de novo' pyrimidine nucleobase biosynthetic process
magnesium ion transport
response to oxidative stress
cell-cell adhesion
autophagy
magnesium ion transmembrane transporter activity
cell adhesion molecule binding
5'-3' RNA exonuclease activity
ribonucleoprotein complex binding
host cell surface receptor binding
protein binding
phosphodiesterase I activity
DNA binding
infected host cell surface knob
extracellular vesicle
external side of plasma membrane
chaperonin-containing T-complex
nucleus
eukaryotic 48S preinitiation complex
nuclear speck
host cell plasma membrane
